# Single cell transcriptomics reveals functionally specialized vascular endothelium in brain

**DOI:** 10.1101/2022.06.10.495613

**Authors:** Hyun-Woo Jeong, Rodrigo Diéguez-Hurtado, Hendrik Arf, Jian Song, Hongryeol Park, Kai Kruse, Lydia Sorokin, Ralf H. Adams

## Abstract

The blood-brain barrier (BBB) limits the entry of leukocytes and potentially harmful substances from the circulation into the central nervous system (CNS). While BBB defects are a hallmark of many neurological disorders, the cellular heterogeneity at the neurovascular interface and the mechanisms governing neuroinflammation are not fully understood. Through single cell RNA sequencing of non-neuronal cell populations of the murine cerebral cortex during development, adulthood, ageing and neuroinflammation, we identify reactive endothelial venules (REVs), a compartment of specialised post-capillary endothelial cells (ECs) that are characterized by consistent expression of cell adhesion molecules, preferential leukocyte transmigration, association with perivascular macrophage populations, and endothelial activation initiating CNS immune responses. Our results provide novel insights into the heterogeneity of the cerebral vasculature and a useful resource for the molecular alterations associated with neuroinflammation and ageing.

## Introduction

The blood-brain barrier (BBB) controls the entry of a wide range of molecules from the circulation into the central nervous system (CNS) and thereby maintains the appropriate chemical and cellular composition of the neuronal ‘milieu’, which is required for the correct function of synapses and neuronal circuits (Zlokovic, 2008, 2010). The BBB also protects the brain against the entry of leukocytes and potentially harmful substances, a function that is compromised in conditions such as multiple sclerosis, cancer, after stroke, or in response to physical brain damage (Arvanitis et al., 2020; Lopes Pinheiro et al., 2016; Thal and Neuhaus, 2014). The neurovascular unit, the anatomical structure underlying the BBB, is comprised of different cell types including endothelial cells, pericytes and astrocytes, which are all located in close proximity and presumably affect each other through reciprocal interactions (Armulik et al., 2011; Liebner et al., 2018).

Previous work has established that leukocyte extravasation – the exit from the blood stream into the tissue – generally occurs in postcapillary venules of the skin, muscle and mesentery, whereas in lung and liver this process is confined to microcapillaries (Strell and Entschladen, 2008). Even in the absence of neuroinflammation, peripherally activated circulating T cells can cross the endothelium of postcapillary vessels and reach the adjacent subarachnoid space. Here, T cells can encounter tissue resident antigen-presenting cells (APCs), but they are unable to traverse the astrocytic basement membrane and glia limitans (Mapunda et al., 2022). In the absence of cognate antigen presented by APCs, T cells undergo apoptosis or reenter the circulation (Mastorakos and McGavern, 2019). Accordingly, immune surveillance and the initiation of CNS immune responses are highly dependent on the cellular and molecular components of the postcapillary venules and the perivascular environment, but we currently lack a comprehensive understanding of the underlying molecular mechanisms.

Transcriptional profiling has recently provided important insights into the cellular composition of the CNS and the arterial-venous zonation of brain vessels during postnatal development and adult homeostasis (Sabbagh et al., 2018; Vanlandewijck et al., 2018; Zeisel et al., 2015). On the other hand, brain ECs demonstrate similar gene expression changes in various BBB dysfunction models, suggesting a common mechanism for compromised BBB function throughout different neurological disorders (Munji et al., 2019). Despite these important insights, the molecular heterogeneity of vascular cells and their dynamic modulation, which are critical for BBB function (Villabona-Rueda et al., 2019), remain insufficiently understood. One critical technical issue is the low abundance of vascular cells relative to neural cell types in the brain.

In this study, we have established a protocol for the depletion of neurons and oligodendrocytes prior to droplet-based single cell transcriptome analysis of non-neuronal cells in mouse cerebral cortex. We have generated a comprehensive resource of vascular gene expression at single cell resolution during postnatal development, adulthood, ageing, and in the demyelinating neuroinflammatory condition of experimental autoimmune encephalomyelitis (EAE), a mouse model of human multiple sclerosis (Ben-Nun et al., 2014). This resulting data provides useful insights into the cellular and molecular heterogeneity of blood vessels in brain and permits identification of functionally specialized reactive post-capillary venules (REVs), which we propose to regulate the infiltration of activated leukocytes during neuroinflammation. Using a computational framework, we have also constructed a detailed cell-to-cell intercommunication map of the brain vasculature.

## Results

### Single cell RNA-sequencing of non-neuronal cell population in mouse cerebral cortex

To enrich vascular and vessel wall-associated cell types while preserving the heterogeneity of these populations, we depleted myelin-associated neurons and oligodendrocytes, the most abundant cell types in CNS, using a bovine serum albumin (BSA)-gradient method (Figure 1-figure supplement 1A, B). Following quality control and data trimming (Figure 1-figure supplement 1C), a total of 29,406 single cells from three different postnatal ages, juvenile (10,796 cells, postnatal day 10), adult (9,871 cells, 7-11 weeks) and aged (8,739 cells, 18 months) were analysed further (Figure 1-figure supplement 2A). Analysis by a nonlinear dimensionality-reduction technique, uniform manifold approximation and projection (UMAP) (Figure 1A-C), and unsupervised hierarchical clustering (Figure 1D, E) established 6 different non-neuronal cell types, namely endothelial cells (EC), microglia (Micro), astroependymal cells (Astro), perivascular macrophages (PVM), mural cells (Mural) and cerebral fibroblast-like cells (Fibro). Each cell type was successfully annotated by known marker genes such as *Flt1* for ECs, *Tmem119* for microglia, *Folr1* for astroependymal cells (including astrocytes, ependymal cells and choroid epithelial cells), *Mrc1* for PVMs, *Rgs5* for mural cells, and *Nov* for the Fibro population (Figure 1E, F). All these markers are expressed specifically and continuously in all age groups analysed (Figure 1-figure supplement 2B-D).

**Figure 1.**
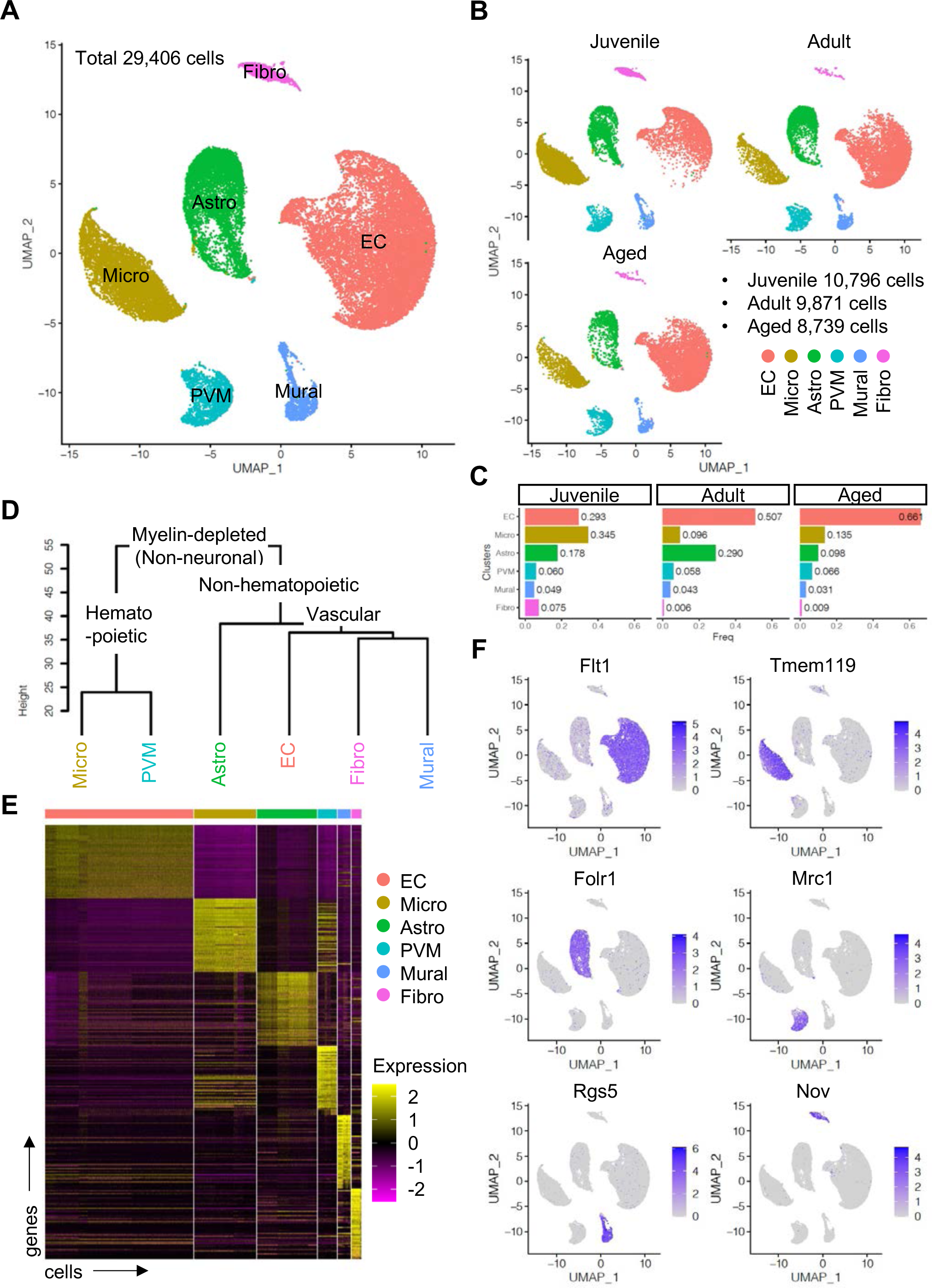
Single cell RNA-seq analysis of non-neuronal cell types in mouse cerebral cortex. **(A)** UMAP plot of 29,406 myelin-depleted single cells from murine cerebral cortex. Colours represent endothelial cells (EC), microglia (Micro), astroependymal cells (Astro), perivascular macrophage (PVM), mural cell (Mural), and cerebral fibroblast (Fibro). (**B**) Split UMAP plots showing separated cells from Juvenile, Adult and Aged samples, respectively. (**C**) Bar plots show the relative frequency of cell types for each age. (**D)** Dendrogram describing the taxonomy of all identified non-neuronal cell types. **(E)** Heatmap indicates the top 50 marker genes for each cell type. **(F)** Expression distribution of the top marker genes for each cell type projected onto the UMAP plot. Colour represents the scaled expression level.

### Unsupervised clustering reveals a diversity of EC subtypes

To gain insight into the heterogeneity of vascular and perivascular cell types, we first analysed ECs during postnatal development using UMAP dimensionality reduction. Juvenile ECs were annotated to 6 different subclusters corresponding to arterial, capillary/pre-capillary arteriolar (CapA), capillary/post-capillary venular (CapV), venous, mitotic, and tip ECs (Figure 2-figure supplement 1A, B). Trajectory analysis indicates that there is one mitotic axis (M) and a tip cell axis (T), which are closely associated with venous populations (V), whereas arterial ECs (A) are found in a different branch of polarization (Figure 2-figure supplement 1C). Distinct transcriptional profiles reveal the expression of venous genes in tip and mitotic ECs (Figure 2-figure supplement 1D, E), while these two populations are also distinguished by specifically expressed transcripts such as *Mcam* for tip cells and *Top2a* for mitotic cells (Figure 2-figure supplement 1F). Pseudotime reconstruction analysis reveals a continuum of venous-to arterial gene expression and also confirmed the presence of venous attributes in tip and mitotic ECs (Figure 2-figure supplement 1G-J), consistent with previous findings that tip and mitotic ECs emerge from veins and are, in part, incorporated into arteries (Pitulescu et al., 2017; Xu et al., 2014).

**Figure 2.**
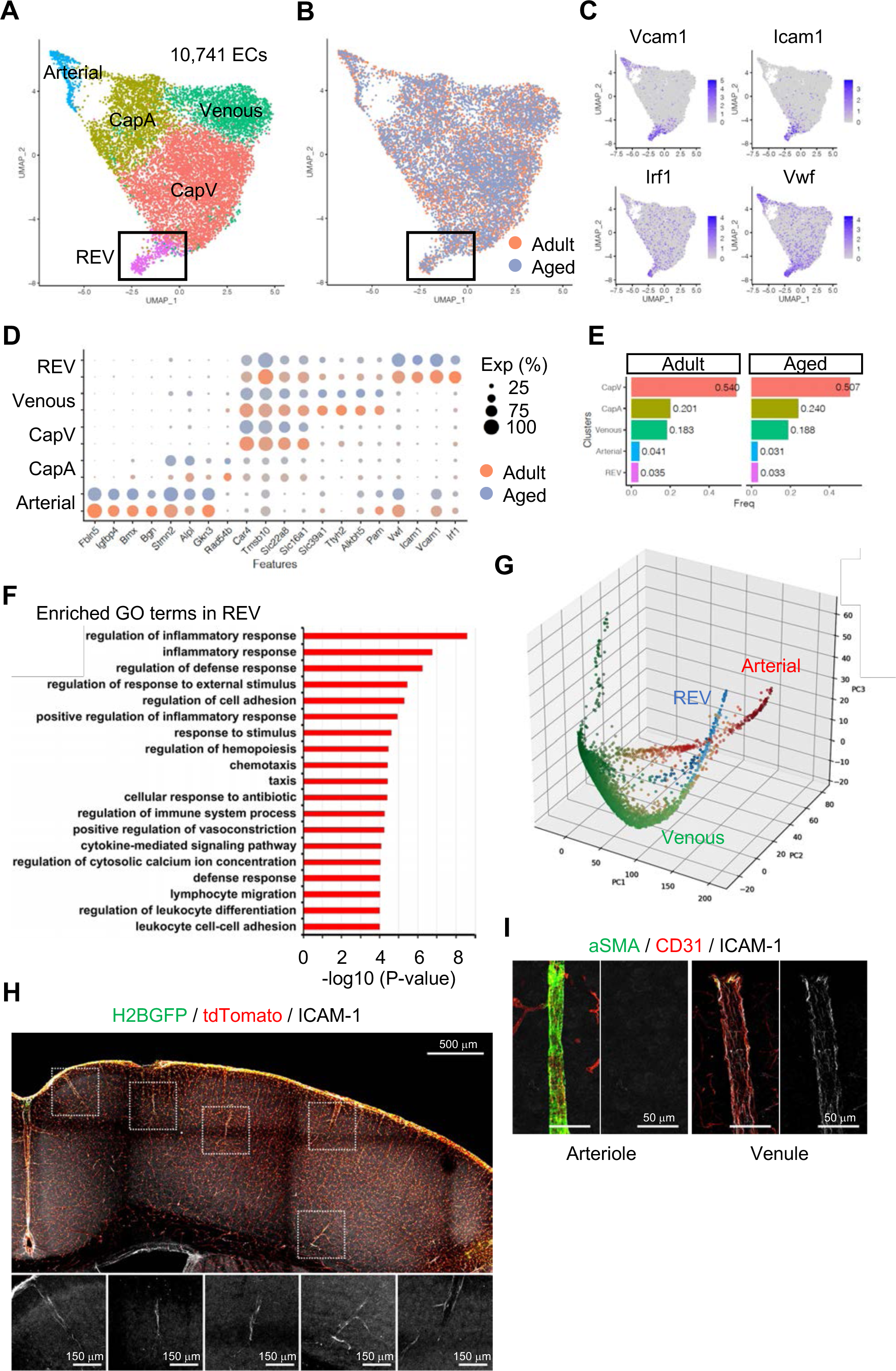
EC sub-clustering and identification of ICAM-1^+^ endothelial population. **(A-B)** UMAP plots of 10,741 adult and aged mouse ECs. Colours represent cell sub-clusters (A) or age groups (B), respectively. Box indicates ICAM-1^+^ reactive endothelial venule (REV) ECs. **(C)** UMAP plots depicting the expression of ICAM-1^+^ EC-enriched genes *Vcam1*, *Icam1*, *Irf1*, and *Vwf*. Colour represents scaled expression level. **(D)** Dot plot showing the expression of top sub-cluster-specific genes, with the dot size representing the percentage of cells expressing the gene and colours representing the average expression of the gene within a cluster. (**E**) Bar plot showing frequency of sub-clusters for adult and aged ECs. **(F)** Top GO biological process terms enriched in REV-specific genes. **(G)** 3D PCA plots generated by MAGIC. Cells are coloured representing the expression of selected subtype marker genes (green: *Alkbh5* and *Tmsb10* for venous ECs; red: *Alpl* and *Fbln5* for arterial ECs; blue: *Icam1* and *Vcam1* for REVs). **(H)** Representative immunofluorescence image for ICAM-1 in adult *Cdh5-H2BGFP/tdTomato* murine brain cortex. Scale bar, 500μm. Panels at the bottom show isolated ICAM-1 signal for each area marked in the overview image, representing different cortical areas. Scale bars, 150μm. **(I)** Immunostaining showing aSMA, CD31 and ICAM-1 expression. Panels on the right show ICAM-1 signal for arteriole and venule, respectively. Scale bars, 50μm.

Next, we analysed adult and aged EC populations. Although there was no striking change in gene expression profiles between adult and aged ECs (Spearman’s correlation coefficient = 0.9852), 407 genes were identified as differentially expressed (Padj < 0.001, Figure 2-figure supplement 2A). Gene set enrichment analysis (GSEA) indicates that genes related to ECM-receptor interaction, focal adhesion and the P53 signalling pathway are enriched in adult ECs, while genes related to cell adhesion molecules, chemokine signalling and T cell receptor signalling are enriched in aged ECs (Figure 2-figure supplement 2B, C), indicating that ECs undergo molecular changes towards a pro-inflammatory state during ageing, which is consistent with recently published work (Chen et al., 2020).

To investigate EC subtypes during homeostasis, we performed subclustering analysis of adult and aged ECs together. As expected, tip and mitotic EC populations are largely absent in these samples, whereas venous, CapV, CapA and arterial ECs are clearly segregated (Figure 2A, B and Figure 2-figure supplement 3A, B). Interestingly, we found one additional minor but distinctive subpopulation of venous ECs (Figure 2A, B; boxed area), which we subsequently termed REV. These ECs are characterized by the expression of *Icam1* and *Vcam1*, encoding intercellular adhesion molecules, and the endothelial activation or dysfunction markers *Vwf* and *Irf1* (Figure 2C, D). It has been reported that endothelial expression of intercellular adhesion molecule-1 (ICAM-1) is essential for transcellular diapedesis (Abadier et al., 2015) but is suppressed by sonic hedgehog (SHH) signalling in the CNS during homeostasis, thereby limiting infiltration of circulating inflammatory leukocytes through the BBB (Alvarez et al., 2011). However, the small subpopulation of venous ECs we found from the single cell transcriptome data shows consistent expression of *Icam1* even in immunologically naïve conditions both in adult and aged mice (Figure 2D, E). To further characterize the *Icam1*-expressing REVs, we performed gene ontology (GO) term enrichment analysis and revealed that 663 genes (Padj < 0.05) predominantly expressed in this cell population were significantly enriched in biological processes involving inflammatory/immune responses, cell adhesion and leukocyte migration and adhesion (Figure 2F). An alternative, nonlinear dimensionality reduction method called Markov affinity-based graph imputation of cells (MAGIC) (van Dijk et al., 2018) also shows a distinct subpopulation of venous ECs with predominant expression of Icam1 and Irf1 (Figure 2G and Figure 2-figure supplement 3C), excluding the possibility that the heterogeneity of cerebral venous ECs is a computational artefact.

Using immunostaining on 100μm-thick brain vibratome sections from transgenic reporter mice expressing nuclear GFP and membrane Tomato specifically in ECs (*Cdh5-H2BGFP/tdTomato*) (Jeong et al., 2017), we confirmed that ICAM-1^+^ REVs are present throughout the cortex of immunologically unchallenged juvenile (Figure 2-figure supplement 3D) and adult mouse brain (Figure 2H). ICAM-1^+^ vessels in the brain parenchyma are predominantly venules with a diameter ranging from 8-50μm (Figure 2-figure supplement 3E) and low to negative coverage by alpha-smooth muscle actin^+^ (αSMA^+^) mural cells (Figure 2I), suggesting that they present a subpopulation of smaller, presumably postcapillary venules where leukocyte extravasation into the brain predominantly occurs. While it was not possible to detect VCAM-1 staining in naïve brains, the cellular frequency of *Icam1* and *Vcam1* double-positive ECs ranges from 1 to 3 % of total ECs in all stages analysed (Figure 2-figure supplement 3F). Postcapillary venules are the main site of leukocyte extravasation in most tissues and in the brain are associated with a perivascular space, defined by the inner endothelial and outer parenchyma or astroglial basement membranes, where activated leukocytes accumulate before entry into the brain parenchyma (Agrawal et al., 2006; Sixt et al., 2001; Wu et al., 2009; Zhang et al., 2020). To test whether the ICAM-1^+^ subpopulation is related to leukocyte infiltration upon an acute inflammation, we intraperitoneally injected adult mice with 10mg/kg of lipopolysaccharides (LPS), perfused the vasculature with ice-cold PBS after 2hr to remove non-adherent cells from the vessel lumen, and performed immunostaining. Interestingly, this revealed the accumulation and penetration of CD3^+^ lymphocytes at ICAM-1^+^ venules prior to significant upregulation of ICAM-1 in other endothelium (Figure 2-figure supplement 3G), suggesting that the ICAM-1^+^ vessel subset might serve as the first entry site of the activated leukocytes to the brain parenchyma in settings of neuroinflammation. Based on these findings, we refer to this vessel compartment as “reactive endothelial venules” (REVs).

### PVMs reside in close proximity of ICAM-1 expressing REV ECs

Brain parenchyma-resident myeloid cell types, namely microglia and PVMs, share expression of myeloid surface markers including CD68, *Fcgr3* (CD16), Cx3cr1 and Aif1 (Figure 3-figure supplement 1A), and are GFP^+^ in *Cx3cr1* reporter mice (Jung et al., 2000) (Figure 3-figure supplement 1B). At the same time, single cell transcriptome analysis shows molecular differences between the two cell populations (Figure 3A), which also correlates with distinct morphologies and localization (Figure 3-figure supplement 1B, C). As the heterogeneity of microglia during development, ageing and brain pathogenesis including Alzheimer’s disease, toxic demyelination and neurodegeneration has been extensively analysed at the single cell level (Hammond et al., 2019; Keren-Shaul et al., 2017; Masuda et al., 2019), we focused on the characterization of PVMs in this study. CD206 is predominantly expressed by PVMs (Figure 3-figure supplement 1D) and CD206^+^ PVMs are localized between the subendothelial basement membrane and GFAP^+^ astrocyte processes (Figure 3B-D). PVMs are clearly distinct from EC-associated pericytes (Figure 3C, asterisks) and *Pdgfra*-expressing adventitial fibroblast-like cells (Figure 3D). Consistent with previous reports showing that PVMs are associated with both arteries and veins in the CNS (Faraco et al., 2017; Faraco et al., 2016), we found PVMs in the proximity of ICAM-1^+^ vessels in cortex, thalamus, hippocampus, midbrain, and floor plate (Figure 3E, F).

**Figure 3.**
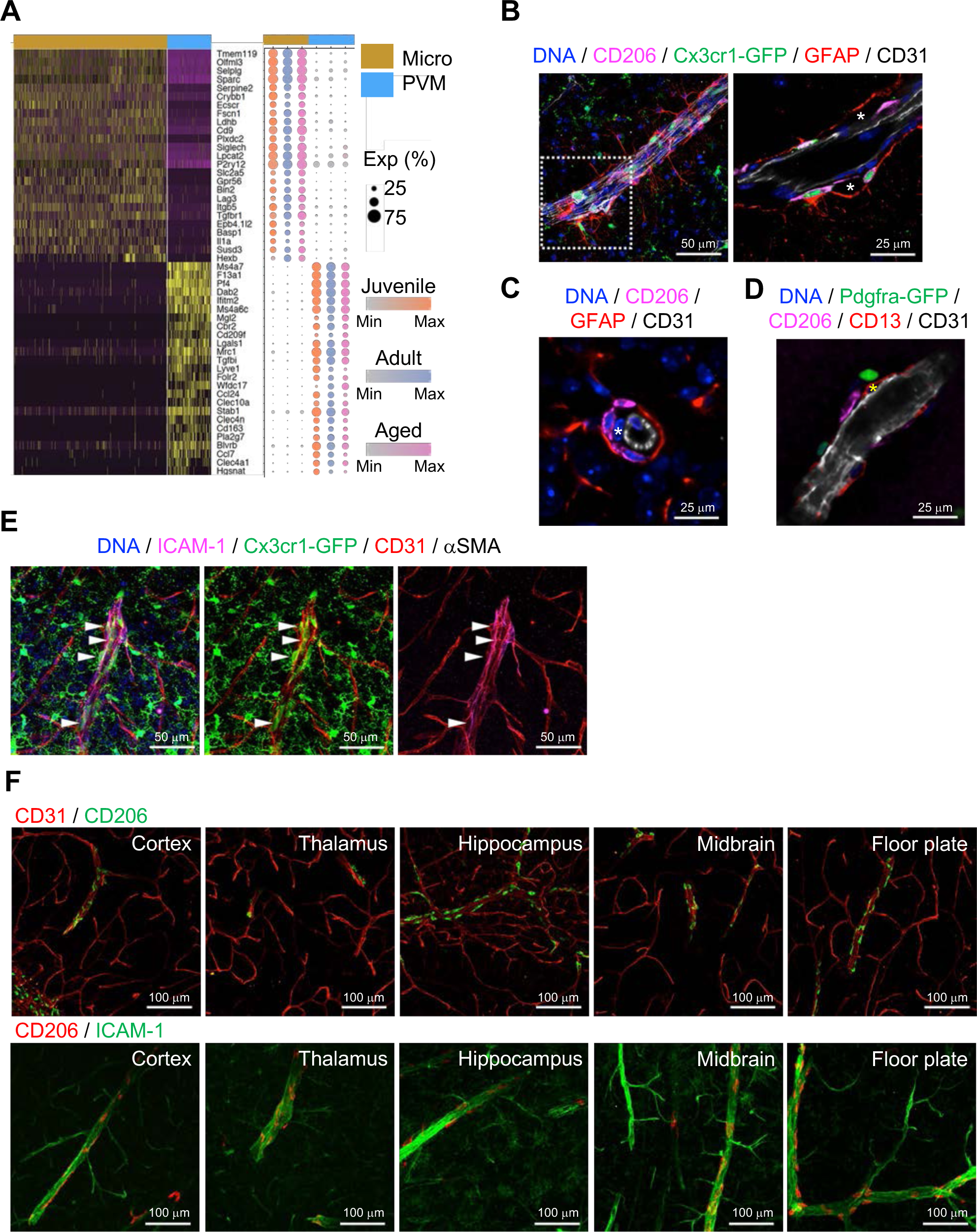
Localization of PVMs. **(A)** Heatmap and expression dot plot for top differentially expressed genes between microglia and PVMs. Dot size represents the percentage of cells expressing each gene and colour keys indicate scaled expression in each age group. **(B)** Immunostaining showing CD206^+^ *Cx3cr1-GFP*^+^ PVMs in perivascular space (asterisk) between ECs (CD31) and astrocyte limitans (GFAP). Scale bars, 50μm (left), 25μm (right). **(C-D)** Spatial arrangement of PVMs (CD206), astrocyte limitans (GFAP), pericytes (CD13, asterisks), peri-vascular fibroblasts (*Pdfgra-GFP*^+^) and ECs (CD31). Scale bars, 25μm. **(E)** Confocal image showing *Cx3cr1-GFP* ^+^ microglia and vessel-associated PVMs (arrowheads) next to ICAM1^+^ αSMA^-^ CD31^+^ REV ECs. Scale bar, 50μm. **(F)** CD206^+^ PVMs are associated with ICAM-1^+^ REVs in the indicated brain regions. Scale bars, 100μm.

Subclustering analysis indicated at least two different PVM populations, namely Lyve1^+^ and MHC class II^+^ (MHCII^+^) PVMs (Figure 4A-C), and differential gene expression analysis revealed unique transcriptomic landscapes of the two subtypes (Figure 4D, Figure 4-figure supplement 1A, B). Immunostaining of Lyve1 and MHCII also shows the two subtypes in association with ICAM-1^+^ vessels (Figure 4E). Interestingly, GSEA for KEGG signalling pathways indicates that Lyve1^+^ PVMs show enriched expression of genes associated with lysosome activity and endocytosis as well as Wnt signalling, whereas MHCII^+^ PVMs show enrichment of terms such as antigen processing and presentation, cell adhesion molecules and Toll-like receptor signalling pathway (Figure 4F, Figure 4-figure supplement 1C). Macrophages can be broadly divided into classical (M1) and alternative (M2) subtypes (Mantovani et al., 2005; Mantovani et al., 2004). M1 polarized macrophages are strongly positive for MHC class II and present antigen to T lymphocytes that neutralize cells with viral or bacterial infections. Macrophages with M2 polarization display homeostatic or anti-inflammatory activity and have a higher endocytic ability compared to M1 macrophages (Tarique et al., 2015). Interestingly, Lyve1^+^ PVMs express anti-inflammatory or immune-suppressive (M2 macrophage-like) polarization genes, such as *Ccr1, Cd163, Cd209a/f, Cd302, Igf1, Il21r, Mrc1, Stab1, Tgfb1*, and *Tslp*, while MHCII^+^ PVMs show increased levels of proinflammatory (M1-like) genes, *Cxcl9, Cxcl10, Cxcl13, Cxcl16, Irf5, Il1a/b, Cxcr4, Il2ra*, and *Tlr2* (Figure 4G). Both PVM subtypes are found at all stages investigated, but the frequency of MHCII^+^ PVMs increases with ageing, which correlates with a decrease in the frequency of Lyve1^+^ PVMs (Figure 4H). Not only the cell population but also the expression levels of MHCII genes are upregulated, whereas *Lyve1* or *Ccl24* expression are downregulated in aged PVMs (Figure 4I). These results indicate that specific PVM subtypes, characterized by distinct immunological signatures, are associated with ICAM-1^+^ REVs.

**Figure 4.**
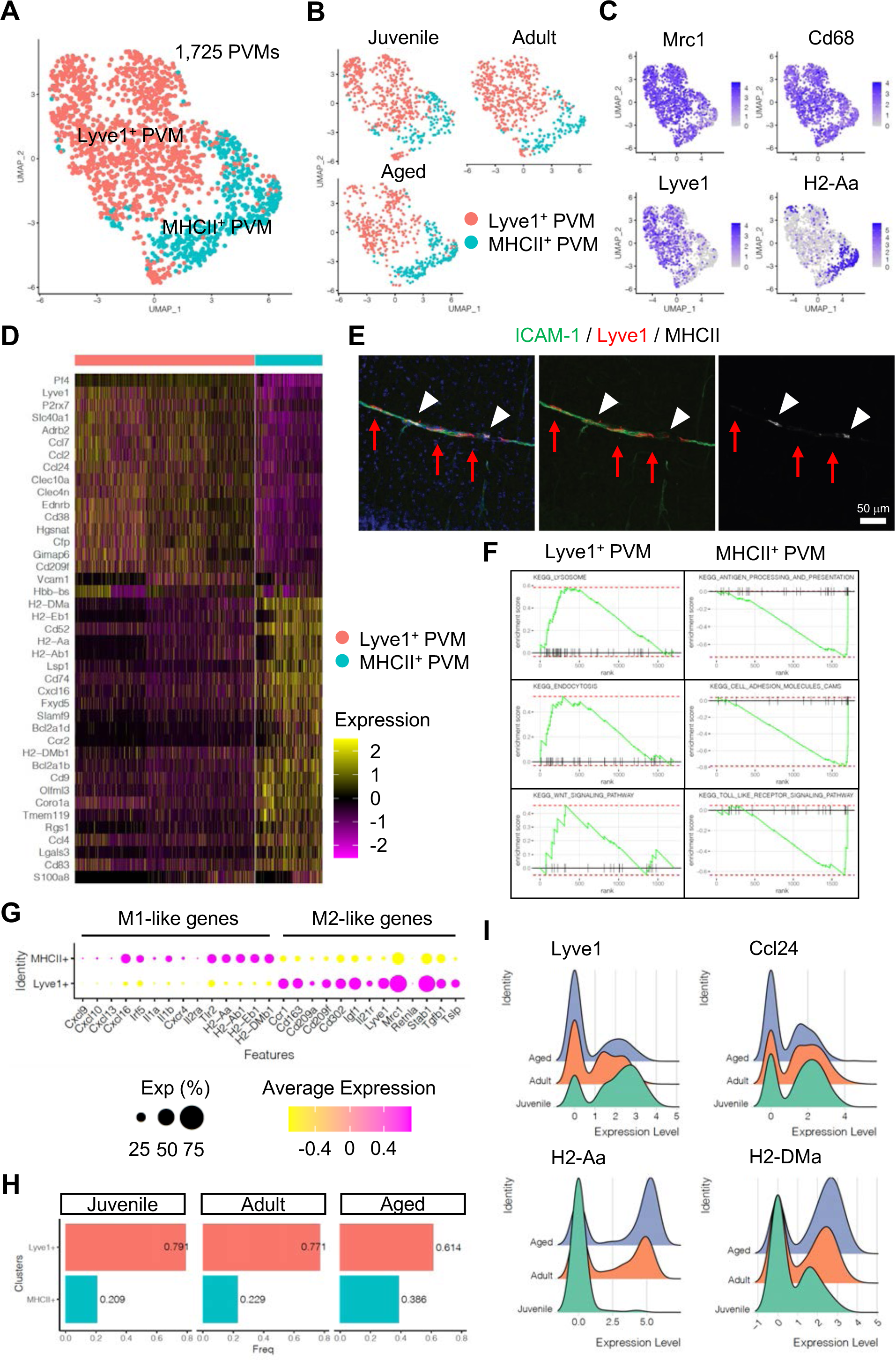
PVM heterogeneity. **(A)** UMAP plot of 1,725 PVMs from juvenile, adult and aged mice distributed into Lyve1^+^ and MHCII^+^ sub-clusters. **(B)** Split UMAP plots of PVMs from Juvenile, Adult and Aged samples. **(C)** UMAP plots depicting the expression of *Mrc1*, *Cd68*, *Lyve1* and *H2-Aa*. Colour represents the scaled expression level. **(D)** Heatmap of top marker genes for Lyve1^+^ and MHCII^+^ PVMs. **(E)** Immunostaining showing Lyve1^+^ (red arrows) or MHCII^+^ PVMs (arrow heads) associated to ICAM-1^+^ REVs. Scale bar, 50μm. **(F)** Representative Gene set enrichment analysis (GSEA) plots for overrepresented KEGG pathways in Lyve1^+^ and MHCII^+^ PVMs. **(G)** Dot plot of genes related to M1 or M2-like phenotypes in PVM subtypes. Dot size represents percentage of cells expressing the gene and colours represents the average expression of each gene. **(H)** Bar plot showing the frequency of PVM subtypes in different age groups. **(I)** Ridge plots of *Lyve1*, *Ccl24* and MHCII genes in PVMs at different ages.

### ICAM-1^+^ ECs are the most reactive EC population in EAE

To extend our analysis of brain ECs and PVMs to neuroinflammatory disease, we analysed the transcriptome of non-neuronal cell types in the brain cortex during experimental autoimmune encephalomyelitis at single cell resolution. In contrast to age-matched control mice, brain-infiltrating inflammatory cells, activated macrophages, neutrophils and lymphocytes were significantly enriched in EAE (Figure 5-figure supplement 1A, B). EAE mice show onset of disease symptoms between 9 and 12 days and peak disease severity between 14 and 18 days after immunization with significant reduction of myelin basic protein (MBP) levels in the brain cortex (Figure 5-figure supplement 1C). The comparative analysis of ECs from healthy adults vs. peak EAE indicates the emergence of inflammatory ECs upon neuroinflammation (Figure 5A, Figure 5-figure supplement 2A). These ECs share the molecular features of CapV (Figure 5-figure supplement 2B) but show higher level of genes related to immune responses such as *Ctla2*, *Lcn2* and *Mt2* (Figure 5-figure supplement 2C).

**Figure 5.**
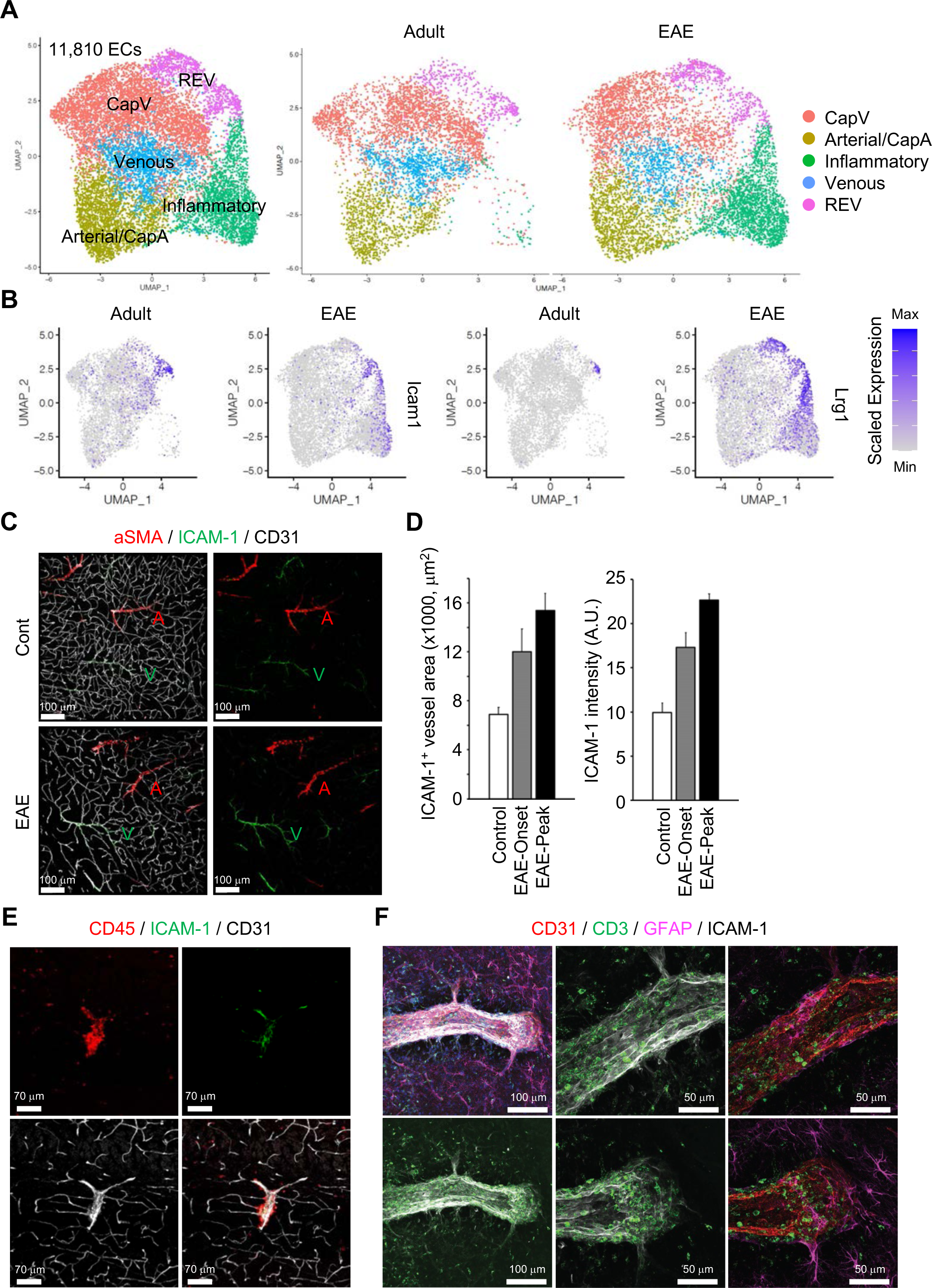
Gene expression changes in ECs by EAE. **(A)** UMAP plots of 11,810 ECs from adult and EAE mice. Colours represent each EC sub-cluster (left) or condition (right). **(B)** UMAP plots depict *Icam1* and *Lrg1* expression in ECs from adult and EAE mice. Colour represents the scaled expression level. **(C)** Immunostaining showing aSMA, CD31 and ICAM-1 expression in control and EAE mice. Arterial (A) and venous vessels (V) are indicated. Scale bars, 100μm. **(D)** Quantification of ICAM-1^+^ vessel area and immunohistochemistry signal intensity in the brain cortex of control and EAE mice at disease onset (EAE Onset) and peak (EAE Peak). Error bars represent mean ± s.e.m. from three animals. **(E)** Immunofluorescence images showing CD45^+^ leukocytes near ICAM-1^+^ REVs in the cerebral cortex of EAE mouse. CD31 indicates all ECs. Scale bars, 70μm. **(F)** High-resolution immunofluorescence images for CD3^+^ lymphocytes, ICAM-1^+^ REVs and GFAP^+^ astrocytes in the cerebral cortex of EAE mouse. CD31 labels all ECs. Scale bar, 100μm (left), 50 μm (middle and right).

REV specific marker genes, *Icam1* and *Lrg1* are also upregulated in inflammatory ECs (Figure 5B). Consistent with published reports (Dopp et al., 1994), both the area and level of endothelial ICAM-1 expression are increased in EAE (Figure 5C, D). Interestingly, ICAM-1 expression in the brain parenchyma persists in REVs and, in peak EAE, expands into adjacent vessels (Figure 5C, Figure 5-figure supplement 3A).

It has been reported that high levels of ICAM-1 expression in ECs promote transcellular diapedesis of encephalitogenic T cells and, in absence of absence of endothelial ICAM-1 and ICAM-2 in mutant mice, EAE symptoms were ameliorated (Abadier et al., 2015). Accordingly, upon EAE, leukocytes accumulate at ICAM-1^+^ REVs and enter the adjacent brain parenchyma (Figure 5E-F, Figure 5-figure supplement 3B).

EAE alters gene expression in REVs (Figure 5-figure supplement 4A, B), including significant upregulation of regulators of blood vessel development, such as *Pdgfb* and *Ctgf*, *Cd74* antigen (invariant polypeptide of the major histocompatibility complex, class II antigen-associated), the hemostasis regulator *Vwf*, an interferon gamma (IFNγ)-stimulated gene mediating host immune responses (*Ch25h*), a gene involved in the cellular stress response (*Maff*), and numerous other genes related to antigen processing and presentation as well as cellular responses to hydrogen peroxide or radiation (Figure 5-figure supplement 4C, D). Collectively, our results indicate that ICAM-1^+^ postcapillary venules or REVs are molecularly specialized and function as a gateway for the entry of activated leukocytes through the BBB.

### PVMs in EAE and cell-to-cell interactome analysis of BBB

The analysis of PVMs in adult and EAE mice indicates that these cells rarely proliferate (Figure 6-figure supplement 1A), and their morphology and localization around ICAM-1^+^ REVs persist in EAE (Figure 6A, B and Figure 6-figure supplement 1B). Nevertheless, EAE induces profound changes in PVM gene expression. MHCII^+^ PVM-enriched genes, such as *Cxcl16*, *Ccl8*, *Cd52*, and MHCII genes, are significantly upregulated in most of the PVMs, whereas Lyve1^+^ PVM-enriched genes, namely *Lyve1*, *Igf1*, *Cd163*, *Ccl24*, *Folr1*, and *Cd209f*, are not changed or decreased (Figure 6D, E). This supports the hypothesis that polarization of macrophages towards inflammatory and non-inflammatory phenotypes is not fixed and that these cells possess plasticity, integrating diverse inflammatory signals with physiological and pathological functions (Jordao et al., 2019; Murray, 2017). In addition, differential gene expression analysis comparing PVMs and brain parenchyma-infiltrated activated macrophages (Mac) further illustrates the molecular differences between PVMs and circulating peripheral monocytes/macrophages (Figure 6F, G), which may prove relevant for potential therapeutic approaches involving the targeting and modulation of different types of macrophages.

**Figure 6.**
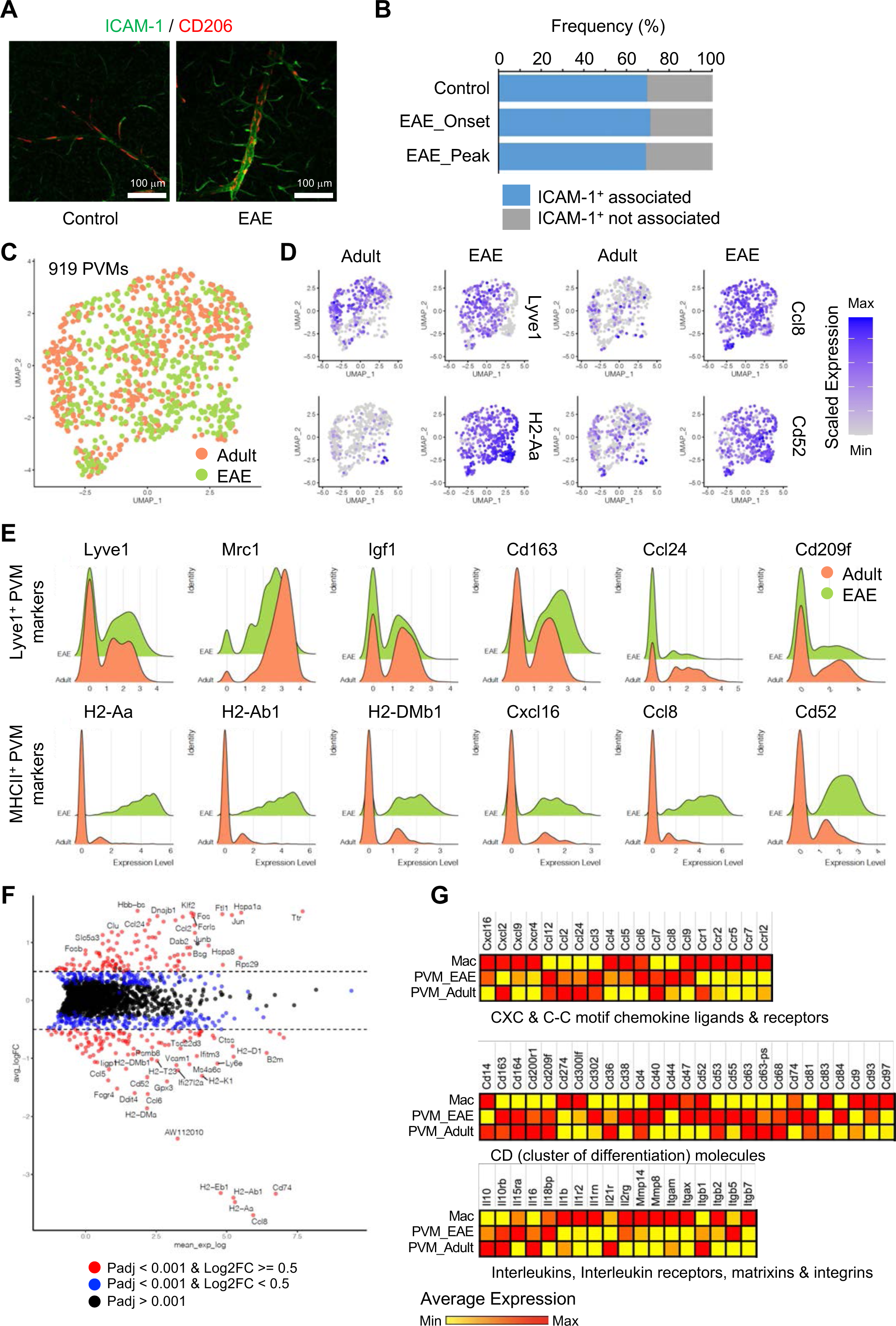
EAE-induced changes in PVMs. **(A)** Representative immunohistochemistry images for CD206^+^ PVMs and ICAM-1^+^ vessels in control and EAE brain cortical parenchyma. Scale bars, 100μm. **(B)** Frequency of CD206^+^ PVMs associated to ICAM-1^+^ vessels in brain cortical parenchyma of control and EAE mice at disease onset (EAE Onset) and peak (EAE Peak). **(C)** UMAP plots of 919 PVMs from adult and EAE mice. **(D)** UMAP plots depicting the expression of *Lyve1*, *H2-Aa*, *Ccl8* and *Cd52* in PVMs of adult control and EAE mice. Colour indicates the scaled expression. **(E)** Ridge plots of Lyve1^+^ and MHCII^+^ PVM marker genes in adult and EAE PVMs. **(F)** MA plot of differentially expressed genes between PVMs from healthy adult and EAE brain. Blue dots, p-adjusted value < 0.001; Red dots, p-adjusted value < 0.001 and Log2 fold change > 0.5. **(G)** Heatmap of selected differentially expressed genes (Padj < 0.001) across adult and EAE PVMs and activated macrophages (Mac). Colour indicates the scaled average expression.

We next used CellPhoneDB (Vento-Tormo et al., 2018) to identify potential receptor-ligand complexes mediating cell-to-cell communication between different cell populations at the BBB. First, we analysed the communication between REVs and adjacent CapV and venous ECs. REVs show expression of various autologous signalling molecules regulating fundamental aspects of EC behaviour, including *Wnt5a*, *Tnfsf12*, *Tgfb1*, *Jag1*, collagens, and immune-regulatory genes with their corresponding receptors (Figure 7A). CapV and venous ECs express higher level of *Esam*, which encodes endothelial cell-selective adhesion molecule, an immunoglobulin-like transmembrane protein associated with endothelial tight junctions. CapV and venous ECs also express the chemokine *Cxcl12* and REVs express the corresponding receptor *Ackr3* encoding CXCR-7, which might enable communication between EC subsets. Potential molecular interactions that upregulated in REVs by EAE include members of the tumor necrosis factor receptor superfamily (TNFRSF), TGFβ receptors, integrins, the HLA class II histocompatibility antigen gamma chain CD74, the receptor tyrosine kinase TEK, and the PLXNB2 receptor for semaphorin ligands (Figure 7A, B). These results also suggest that REVs are a major signal distributor among the EC populations in CNS, which is further enhanced by EAE (Figure 7C). We also identify putative signalling interactions between REVs and other cell types in the brain vasculature during homeostasis and neuroinflammation (Figure 7-figure supplement 1). Notably, REVs express growth factors including PDGF-B, CTGF, and TGFβ;1, which regulate cellular behaviour, tissue remodelling and angiogenesis. Signalling to other cell types is further enhanced by EAE, which involves the activation of various signals in astroependymal cells. By contrast, expression of PDGF-A, IGF2 and PGF in mural cells and of IGF1, TGFB1 and IGFBP4 in PVMs, for example, was not significantly changed upon EAE (Figure 7D). Our results collectively suggest that postcapillary venules form a specialised vessel compartment in the CNS. We propose that REVs might play roles in the regulation of immune surveillance in the CNS both under homeostatic conditions but also in pathogenic neuroinflammation.

**Figure 7.**
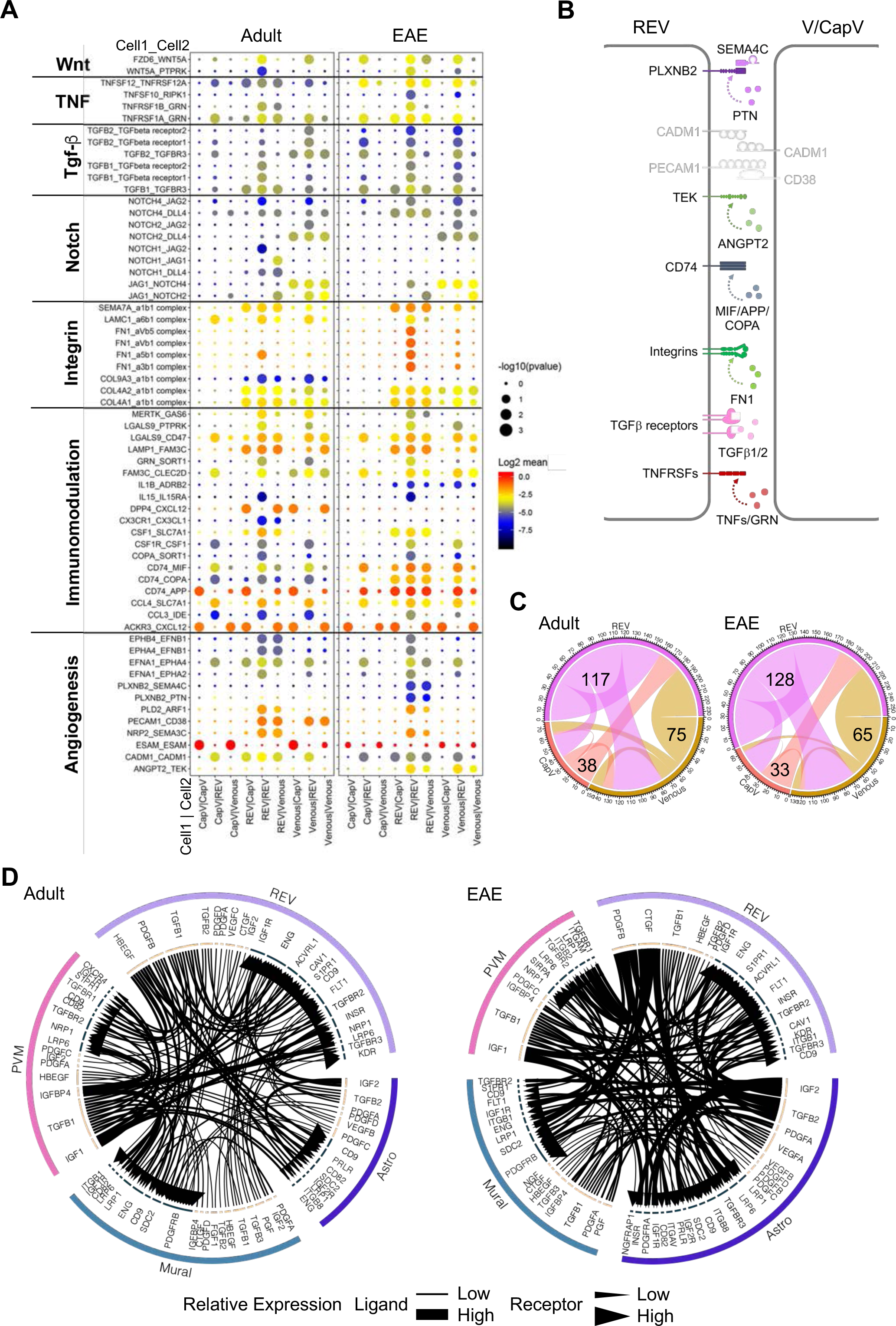
Interactions between different EC subpopulations and BBB cell types. **(A)** Overview of potential ligand-receptor interactions for CapV, REV and Venous EC populations in adult and EAE based on gene expression. Circle size indicates P values. Colour key indicates the means of the average expression levels of interacting molecules. **(B)** Diagram of the main ligand-receptor interactions regulated by EAE. **(C)** Diagram of the numbers of ligand-receptor interactions on CapV, REV and Venous EC populations in healthy adult and EAE brain. **(D)** Circos plots for ligand-receptor interactions between REV ECs, mural cells (Mural), PVMs and astroependymal cells (Astro) in adult and EAE. Each plot shows top 100 highly expressed interactions. The lines and arrow heads are scaled to indicate the relative expression level of the ligand and receptor, respectively.

## Discussion

Here, we provide a comprehensive single cell transcriptomic atlas of non-neural components of the murine brain vasculature during growth, adulthood, ageing, and in EAE. Using this data, we establish the existence of a specialized vessel subtype, which is characterized by a distinct endothelial gene expression profile and presence of intercellular adhesion molecules. In response to pro-inflammatory (LPS) stimulation or in the neuroinflammatory condition of EAE, REVs, but not the far more abundant ICAM-1^-^ vessels, are associated with leukocytes, suggesting that the ICAM-1^+^ subset of postcapillary venules serves as the first site of immune cell recruitment into brain parenchyma. The association of MHCII^+^ and Lyve1^+^ PVM populations with REVs might mediate immunomodulatory responses and the predominant gene expression changes in REVs upon immunological challenge highlight further that these vessels might be critical for the immune privileged status of the CNS with potential implications for ageing and neurodegenerative diseases.

Several previous studies on scRNA-seq analysis of murine or human brain have recovered only a limited number of vascular cells due to the comparably high abundance of neural cells (Han et al., 2020; Zeisel et al., 2015). In other publications, fluorescence-activated cell sorting (FACS) or microbead-mediated isolation of ECs as well as of both ECs and mural cells have provided insight into the organ-specific specialization, transcriptional regulation and arterial-venous zonation of brain vessels in the healthy organism (He et al., 2018; Kalucka et al., 2020; Sabbagh et al., 2018; Vanlandewijck et al., 2018). While sorting based on certain cell surface markers or fluorescent reporters enables the efficient enrichment of the desired cell populations, it has to be considered that these approaches are biased and will miss cells lacking expression of the relevant markers. This limitation is avoided by our demyelination approach, which does not rely on the expression of specific cell surface proteins or fluorescent reporters. Despite of the differences in the experimental approach, data generated by Vanlandewijck et al. (Vanlandewijck et al., 2018) confirms the enriched expression of Icam1 and Vwf in a fraction of venous ECs, which is consistent with our own findings. Transcriptomic changes in brain ECs associated with the aging have been also investigated previously and it has been proposed that soluble brain EC-derived Vcam1, generated by shedding, activates microglia and impairs the function of hippocampal neural precursors (Chen et al., 2020; Yousef et al., 2019). Our own study confirms that aging increases transcripts associated with inflammation in brain ECs, and, in particular, in REVs. Similarly, EAE results in the upregulation of proinflammatory gene expression in ECs with potential implications for neuroinflammation and disease development.

In the healthy organism, the CNS contains a highly selective barrier system to restrict peripheral immune cell entry into brain parenchyma. During pathophysiological conditions, activated T cells can enter the perivascular space independent of antigen specificity (Kawakami et al., 2005) along postcapillary venules (Raine et al., 1990). However, only after acquiring the ability to move across the glia limitans by recognition of cognate antigen displayed by perivascular APCs, T cells traverse into the parenchyma (Lodygin et al., 2013). Firm attachment of T cells to ECs to resist the vascular shear stress and prolonged crawling before diapedesis are also critical for immune cell extravasation (Steiner et al., 2010). Our findings suggest that functionally specialized REV ECs enable the sustained immune surveillance process alongside the postcapillary venules, serving as the cellular gateway of BBB. In this sense, REVs share functional features with high endothelial venules (HEVs) in secondary lymphoid organs that support circulating lymphocyte extravasation and physical contact with APCs (Girard and Springer, 1995). Because of the restricted afferent and efferent communication with lymphatic tissue in CNS parenchyma, these specialized postcapillary ECs are probably relevant for fine-tuned CNS immune reactions.

Our work also shows transcriptionally distinguishable subtypes of brain resident PVMs, physically associated with REVs in postcapillary venules, which might serve as APCs for CNS immune surveillance and thereby as gate-keepers at the BBB. It has been reported that PVMs are derived from early yolk-sac-derived erythromyeloid progenitors, similar to microglia, and have minimal turnover during homeostasis (Goldmann et al., 2016). The transcriptional continuum between the PVM subtypes and the coherent activation upon EAE reflect the ability of PVMs to swiftly adapt to changing immunological and environmental cues. Nevertheless, PVMs retain distinct molecular characteristics relative to infiltrating peripheral macrophages even though all these cells are exposed to the same perivascular microenvironment. More detailed understanding of the mechanisms that determine the behaviour of PVMs and peripheral macrophages upon neuroinflammation will be important, especially for attempts to target specific cell populations and modulate local immune responses. Delivery of drugs or therapeutic cells (e.g. chimeric antigen receptor T cells) across the BBB is a limiting factor in the future development of new therapeutics for the brain diseases, such as Alzheimer’s disease and various brain tumours (Pardridge, 2019).

The sum of our work reveals critical aspects of vascular heterogeneity in the CNS and thereby provides provides a valuable resource for cell-cell interactions and the molecular modulation of immune cell trafficking into the CNS, with relevance for brain function in health and disease.

## Methods

### Mice

C57BL/6 female mice were used unless stated otherwise. *Cdh5-H2BGFP*/*tdTomato* (Jeong et al., 2017)*, Cx3cr1-GFP* (Jung et al., 2000) and *Pdgfra-GFP* transgenic mice were used to specifically label EC, Micro + PVM and Fibro, respectively. All animal experiments were performed in compliance with the relevant laws and institutional guidelines, approved by local animal ethics committees, and conducted with permissions granted by the Landesamt für Natur, Umwelt und Verbraucherschutz (LANUV) of North Rhine-Westphalia, Germany.

### EAE and LPS treatment

EAE was induced as previously described (Gerwien et al., 2016). In brief, 150 μl of 0.75mg/ml myelin oligodendrocyte glycoprotein 33 to 55 peptides (MOG35–55) mixed with Complete Freund’s Adjuvant (FS8810; Sigma-Aldrich) was injected subcutaneously into the tail base of 7 to 11-week-old female C57Bl/6 mice. On the day of immunization and on day 2, 100 μl of 2μg/ml pertussis toxin (P7208; Sigma-Aldrich) was injected intravenously. Mice were monitored daily for development of disease symptoms. EAE was graded on a 0 to 5 scale as follows: score 0, no clinical symptoms; score 1, flaccid tail; score 2, hindlimb weakness or partial paralysis; score 3, severe hindlimb weakness or paralysis; score 4, forelimb paralysis; and score 5, paralysis in all limbs or death. Mice typically reached a peak of disease symptoms between days 14 and 18 after immunization, before disease resolution. The disease typically correlates with weight loss; killing of mice was required when weight loss exceeded 20 to 30% of the initial body weight. For acute LPS treatment, mice were intraperitoneally injected with a single dose of 10mg/kg lipopolysaccharides (L2630-25MG; Sigma-Aldrich). Control animals received the same volume of PBS (data not shown). Mice were sacrificed at 2 hours after the injection.

### Single cell RNA-Seq library preparation and sequencing

Mice were perfused transcardially with ice-cold PBS under anaesthesia. Brains were isolated and dabbed with filter paper for the removal of meninges. After the removal of olfactory bulbs and cerebellum, brain cortex tissues were dissected and digested using the enzyme-cocktail solution of 30U/ml papain (LK003153; Worthington) and 0.125mg/ml Liberase TM (5401119001; Sigma-Aldrich) in DMEM (Sigma-Aldrich) for 60 min at 37 °C. Homogenized tissues were combined with 1.7-fold volume of 22% bovine serum albumin in PBS and then centrifuged at 1,000x g for 10 min for myelin removal. After washing with ice-cold DMEM, dead cells and debris were removed using ClioCell nanoparticles (Amsbio) according to the manufacturer’s instructions. Single cells were counted using Luna-II automated cell counter (Logos Biosystems) and captured using the 10X Chromium system (10X Genomics). We pooled tissues from three (for aged and EAE samples) or six (for juvenile and adult samples) mice to reduce the effect of sample-dependent individual variations. Libraries were prepared according to the manufacturer’s instructions using Chromium Single Cell 3’ Library & Gel Bead Kit v2 (10X Genomics) and sequenced on the Illumina NextSeq 500 using High Output Kit v2.5 (150 cycles, Illumina) for 26 bp + 98 bp paired-end reads with 8bp single index aiming raw sequencing depth of >20,000 reads per cell for each sample.

### Single cell RNA-seq data analysis

Sequencing data were processed with UMI-tools(Smith et al., 2017) (version 1.0.0), aligned to the mouse reference genome (mm10) with STAR (Dobin et al., 2013) (version 2.7.1a), and quantified with Subread featureCounts (Liao et al., 2014) (version 1.6.4). Data normalization, detailed analysis and visualization were performed using Seurat package (Butler et al., 2018) (version 3.1.5). For initial quality control of the extracted gene-cell matrices, we filtered cells with parameters nFeature_RNA > 500 & nFeature_RNA < 6000 for number of genes per cell and percent.mito < 25 for percentage of mitochondrial genes and genes with parameter min.cell =3. Filtered matrices were normalized by LogNormalize method with scale factor=10,000. Variable genes were found by FindVariableFeatures function with parameters of selection.method = “vst”, nfeatures = 2000, trimmed for the genes related to cell cycle (GO:0007049) and then used for principal component analysis. FindIntegrationAnchors and IntegrateData with default options were used for the data integration. Statistically significant principal components were determined by JackStraw method and the first 12 principal components were used for UMAP non-linear dimensional reduction. Unsupervised hierarchical clustering analysis was performed using FindClusters function in Seurat package. We tested different resolutions between 0.1 ∼ 0.9 and selected the final resolution using clustree R package to decide the most stable as well as the most relevant for our previous knowledges. Cellular identity of each cluster was determined by finding cluster-specific marker genes using FindAllMarkers function with minimum fraction of cells expressing the gene over 25% (min.pct=0.25), comparing those markers to known cell type-specific genes from previous studies.

Data were further trimmed for clusters of multiplets, low quality cells (mitochondrial gene-enriched), contaminated neurons (Tubb3^+^), oligodendrocytes (Mbp^+^), lymphocytes (Igkc^+^ or Nkg7^+^), red blood cells (Hba−a1^+^) and ependymal cells (Tmem212^+^) and then reanalysed. For subclustering analysis, we isolated specific cluster(s) using subset function, extracted data matrix from the Seurat objectusing GetAssayData function and repeated the whole analysis pipeline from data normalization.

Differentially expressed genes (DEGs) were identified using the non-parametric Wilcoxon rank sum test by FindMarkers function of Seurat package. We used default options for the analysis if not specified otherwise. Results were visualized using EnhancedVolcanoR package (version 1.10.0). FeaturePlot, VlnPlot and DotPlot functions of Seurat package were used for visualization of selected genes. The “VlnPlot” function of Seurat package was used for violin plots to show the expression level of selected genes with log normalized value by default.

Monocle (Trapnell et al., 2014) (version 2.12.0) was used for pseudotime trajectory analysis. We imported Seurat objects to Monocle R package and then performed dimensionality reduction with DDRTree method with parameters max_components=2 and norm_method=”log”. Cell cycle phases were classified by cyclone function of scran (Lun et al., 2016) (version 1.14.5). We also used MAGIC (van Dijk et al., 2018) (version 1.4.0.9000) for an alternative dimensionality reduction and CellPhoneDB (Vento-Tormo et al., 2018) for ligand-receptor interactome analysis. An R package iTALK (https://doi.org/10.1101/507871) was used for the ligand-receptor interactome analysis. For the analysis of each single sample, top 50% of genes in their mean expression values were selected and used for ligand-receptor pair identification using FindLR function with datatype = mean count. All the analysis scripts are available from vignettes of original software webpage of Seurat, Monocle, MAGIC, CellPhoneDB or iTALK. No custom code or mathematical algorithm other than variable assignment was used in this study.

### Flow cytometry

Brain cortex tissues from transcardially perfused P10 *Cdh5-H2BGFP*/*tdTomato* transgenic mice were dissected and treated by different digestion conditions, namely papain solution, papain + liberase cocktail solution, or papain + liberase cocktail solution followed by myelin removal. EC-specific H2BGFP and tdTomato fluorescences were directly analysed after single / live cell gating in forward scattered / side scattered plot using FACSVerse flow cytometer (BD). FlowJo software (version 10.5.3, FlowJo, LLC) was used for further analysis.

### Immunohistochemistry

Mice were perfused transcardially with ice-cold PBS and subsequently with 2% paraformaldehyde (PFA) under anaesthesia. Whole brain tissues were dissected and further fixed with 4% PFA or 100% methanol at 4°C for overnight. Methanol-fixed samples were rehydrated by serial incubation (15-20 min each, at RT) in increasing concentrations of PBS:methanol solution (25, 50, 75 and 100% PBS). The fixed brains were glued to a mounting block with cyanoacrylate glue (48700; UHU), submerged in ice-cold PBS and sliced with 100µm-thickness using vibrating blade microtome (VT1200, Leica).

Sections were blocked and permeablilized by 1% BSA and 0.5% Triton X-100 in PBS for overnight. Incubation with blocking/permeabilization solution containing primary antibodies at 4 °C for overnight was followed by secondary antibody staining using suitable species-specific Alexa Fluor-coupled antibodies (Invitrogen) and flat-mounting in microscope glass slides with Fluoromount-G (0100-01; SouthernBiotech). The following primary antibodies were used for immunostaining: rabbit anti-ICAM-1 (Abcam ab222736, 1:100), rat anti-ICAM-1 (BioLegend 116102, 1:100), mouse anti-αSMA-Cy3 (Sigma C6198, 1:300), mouse anti-αSMA-660 (eBioscience 50-9760-82, 1:300), chicken anti-GFP (2BScientific Ltd. GFP-1010, 1:300), goat anti CD31 (R&D Systems AF3628, 1:200), rabbit anti-CD206 (Abcam ab64693, 1:100), rat anti-CD206 (BioRad MCA2235T, 1:100), rabbit anti-GFAP (DAO Z0334, 1:200), rat anti-CD13 (AbD Serotec MCA2183GA, 1:100), hamster anti-CD3e-FITC (eBiosciences 11-0031, 1:100), rabbit anti ColIV (AbD Serotec 2150-1470, 1:100), rat anti-CD68 (Abcam ab53444, 1:200), goat anti-Olig2 (R&D Systems AF2418, 1:100), rabbit anti-MHC Class II (Abcam ab180779, 1:100), rat anti-CD45 (Becton Dickinson 550539, 1:200) and rat anti myelin basic protein (MBP, Abcam ab7349, 1:200). The following donkey-raised secondary antibodies (all in 1:400 dilution unless otherwise stated) were used for immunostaining: anti-rabbit IgG conjugated to Alexa Fluor (AF) 488 (Thermo Fisher Scientific A21206), anti-chicken IgY AF488 (Jackson ImmunoResearch 703-545-155), anti-goat IgG AF488 (Invitrogen, A-11055), anti-rat IgG Cy3 (Jackson ImmunoResearch 712-165-153), anti-rabbit IgG AF546 (Thermo Fisher Scientific A10040), anti-rat IgG AF594 (Thermo Fisher A21209), anti-rabbit IgG AF594 (Thermo Fisher Scientific A21207), anti-rabbit IgG AF647 (Thermo Fisher Scientific A31573), and anti-goat IgG AF647 (Thermo Fisher Scientific A21447). Streptavidin AF405 (Invitrogen S32351, 1:200) was used for detection of biotinylated-IB4 stained samples. Nuclei were counterstained with 4’,6-diamidino-2-phenylindole (DAPI, Sigma, D9542) diluted at 1 μg ml^-1^ together with the secondary antibodies.

### Statistics and reproducibility

No statistical methods were used to predetermine sample size. The experiments were not randomized and investigators were not blinded to allocate during experiments and outcome assessment.

Data sets with normal distributions were analysed with unpaired Student’s two-tailed t-tests to compare two conditions. Results are depicted as mean ± s.e.m. as indicated in figure legends. All experiments for quantitative analysis and representative images were reproduced at least three times.

### Data availability

Raw data (fastq files) and processed data (gene counts) for single cell RNA-Seq analysis have been deposited in the Gene Expression Omnibus with the primary accession number, GSE133283.

## Author contributions

R.H.A. supervised the project. H.-W.J. conceived the project and performed all experiments and analyses. R.H.-D., H.P. and H.A. performed immunohistochemistry. J.S. and L.S. performed active EAE induction and contributed to conceptual aspects of the manuscript. K.K. produced web database. H.-W.J. and R.H.A. wrote the manuscript.

## Acknowledgments

This work was supported by the Max Planck Society, the University of Münster and the German Research Foundation (DFG) CRC1366A01, CRC1009A02, SO285/11-1, and the Cluster of Excellence “Cells in Motion” (EXC1003).

---

**Figure 1-figure supplement 1.**
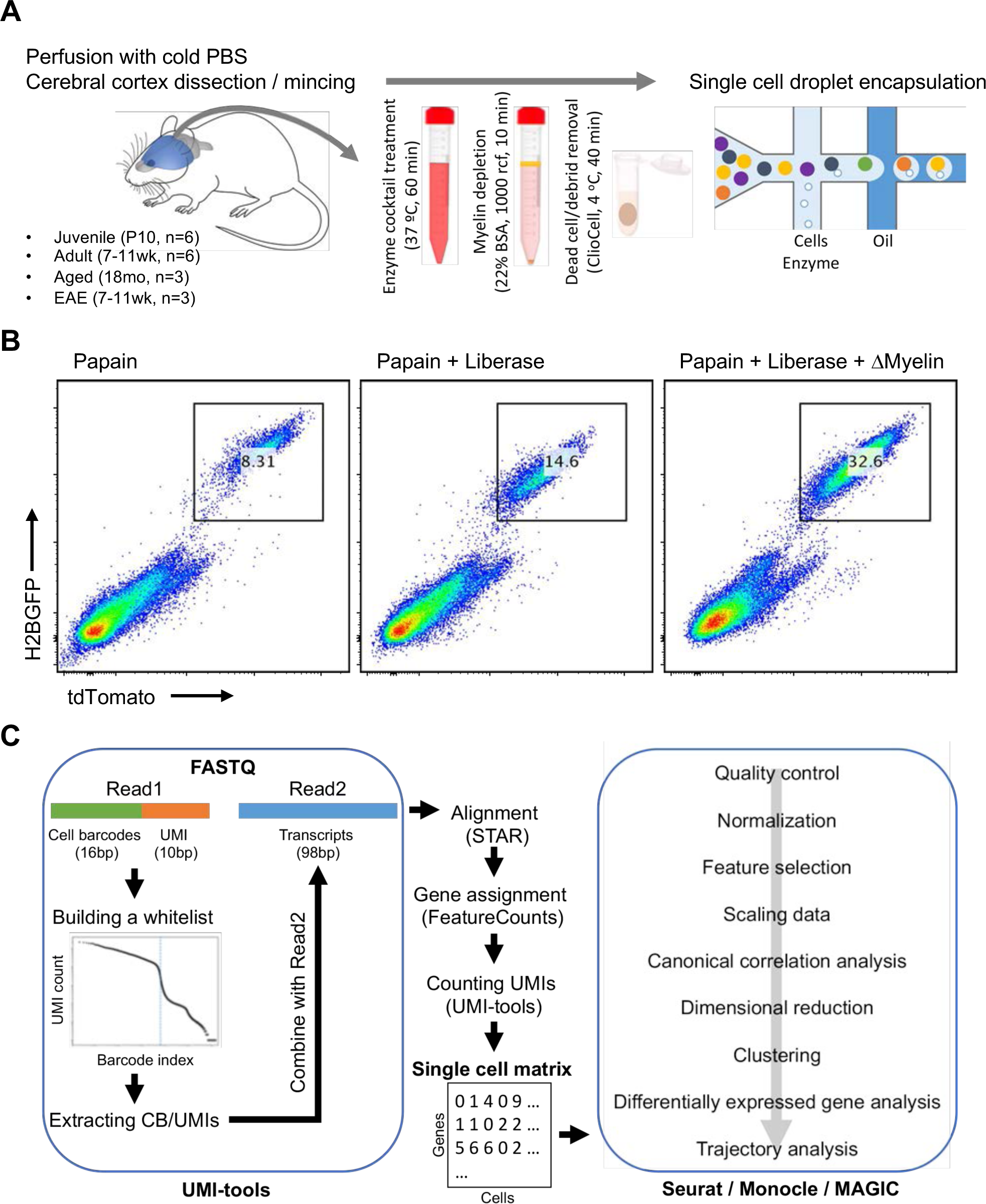
Analysis of vascular and perivascular cell types in mouse cerebral cortex using single cell RNA-seq. **(A)** Workflow for the enrichment of blood vessel-associated cell types from cerebral cortex of juvenile (P10), adult (7-11 weeks), aged (18 months) and EAE (7-11 weeks, peak of disease) mice. **(B)** Flow cytometry plots depicting enrichment of ECs from cerebral cortex of P10 *Cdh5-H2BGFP/tdTomato* transgenic mice using different tissue dissociation methods. **(C)** Pipeline for the single cell RNA-seq data analysis.

**Figure 1-figure supplement 2.**
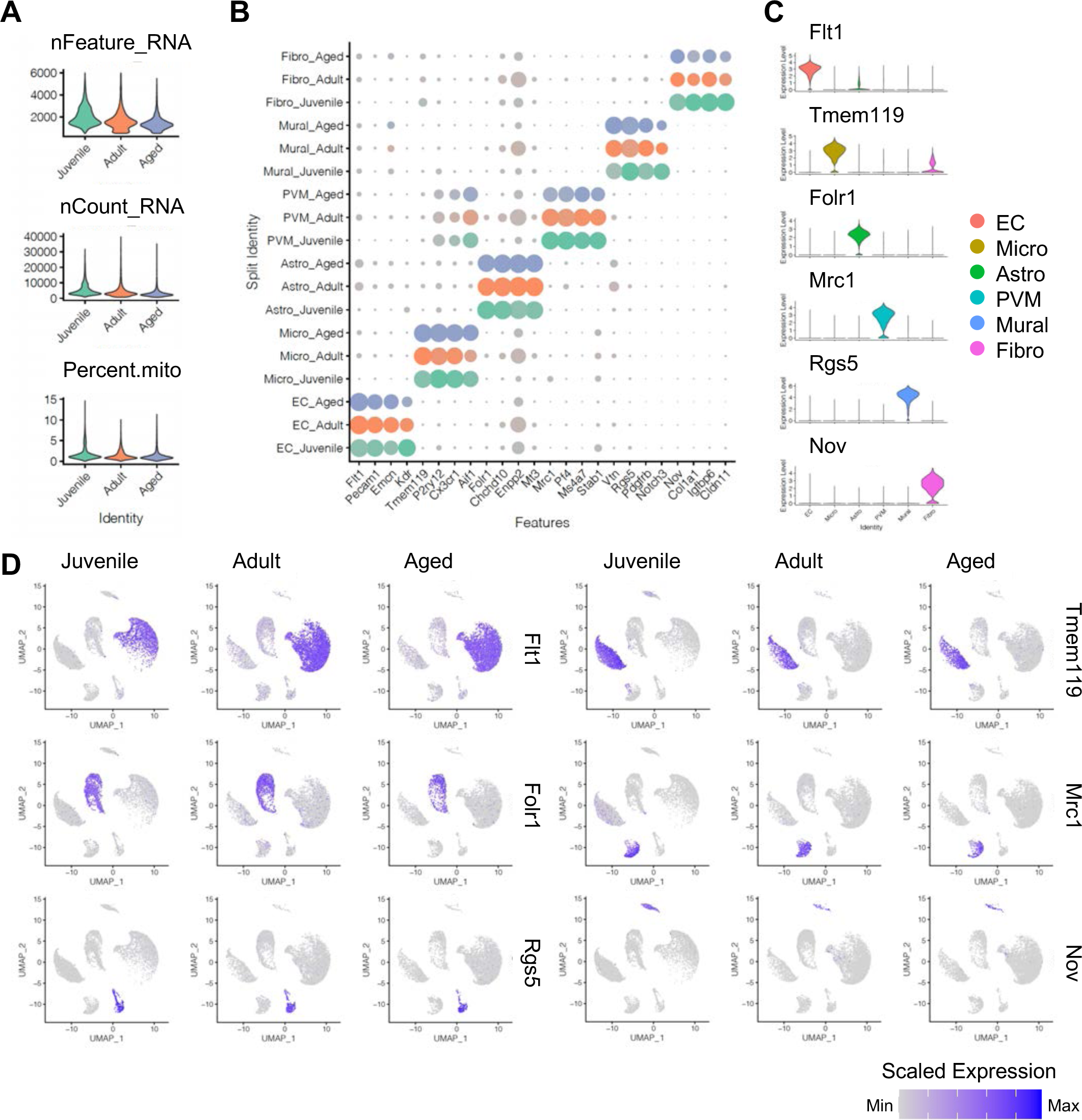
Expression of major cell type marker genes. **(A)** Violin plots displaying number of genes (nFeature_Gene), number of unique molecular identifiers (UMI; nFeature_UMI) and percentage of mitochondrial genes (percent.mito) in juvenile, adult and aged samples. **(B)** Dot plot showing the expression of top cell type-specific genes, with the dot size representing the percentage of cells expressing the gene and colours representing the average expression of the gene within a cluster. **(C-D)** The top cell type-specific genes are shown in violin plots (C) and UMAP plots (D). Colour represents the scaled expression level.

**Figure 2-figure supplement 1.**
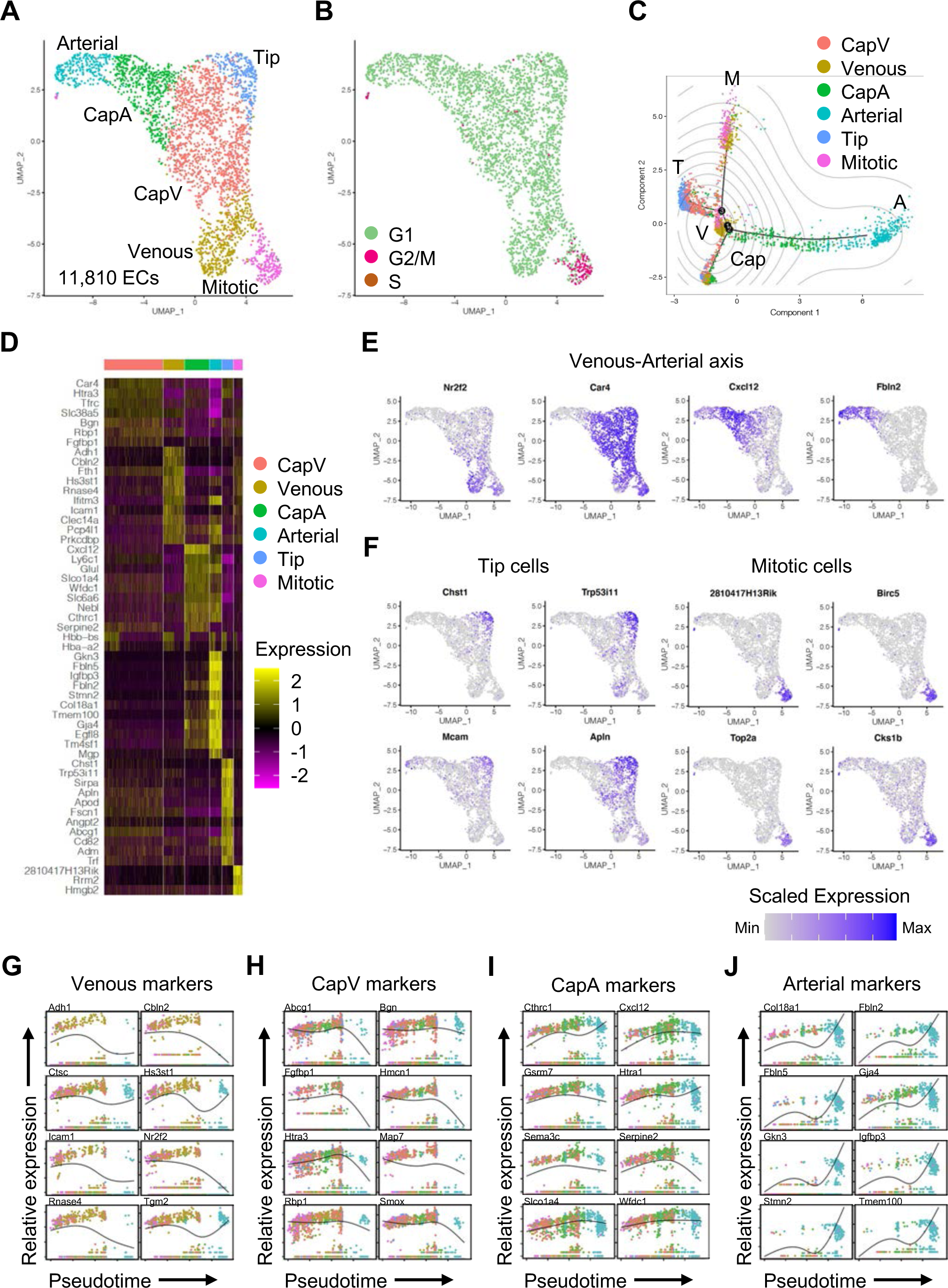
Subclustering analysis of ECs during postnatal development. **(A-B)** UMAP plots of juvenile ECs. Colours represent EC sub-clusters (A) or cell cycle phases (B). **(C)** Trajectory analysis of juvenile ECs depicting centrally located venous ECs (V) and polarized differentiation paths of mitotic (M), tip (T), capillary (Cap) and arterial (A) ECs. Contour plots indicate two-dimensional Gaussian kernel density estimates and colours represent each sub-cluster. **(D)** Heatmap of the top 12 marker genes for each sub-cluster of juvenile ECs. **(E-F)** UMAP plots of marker genes for each sub-cluster in juvenile ECs. **(G-J)** Kinetics plots showing relative expression of venous to arterial marker genes across pseudotime. Colours represent sub-clusters shown in (A).

**Figure 2-figure supplement 2.**
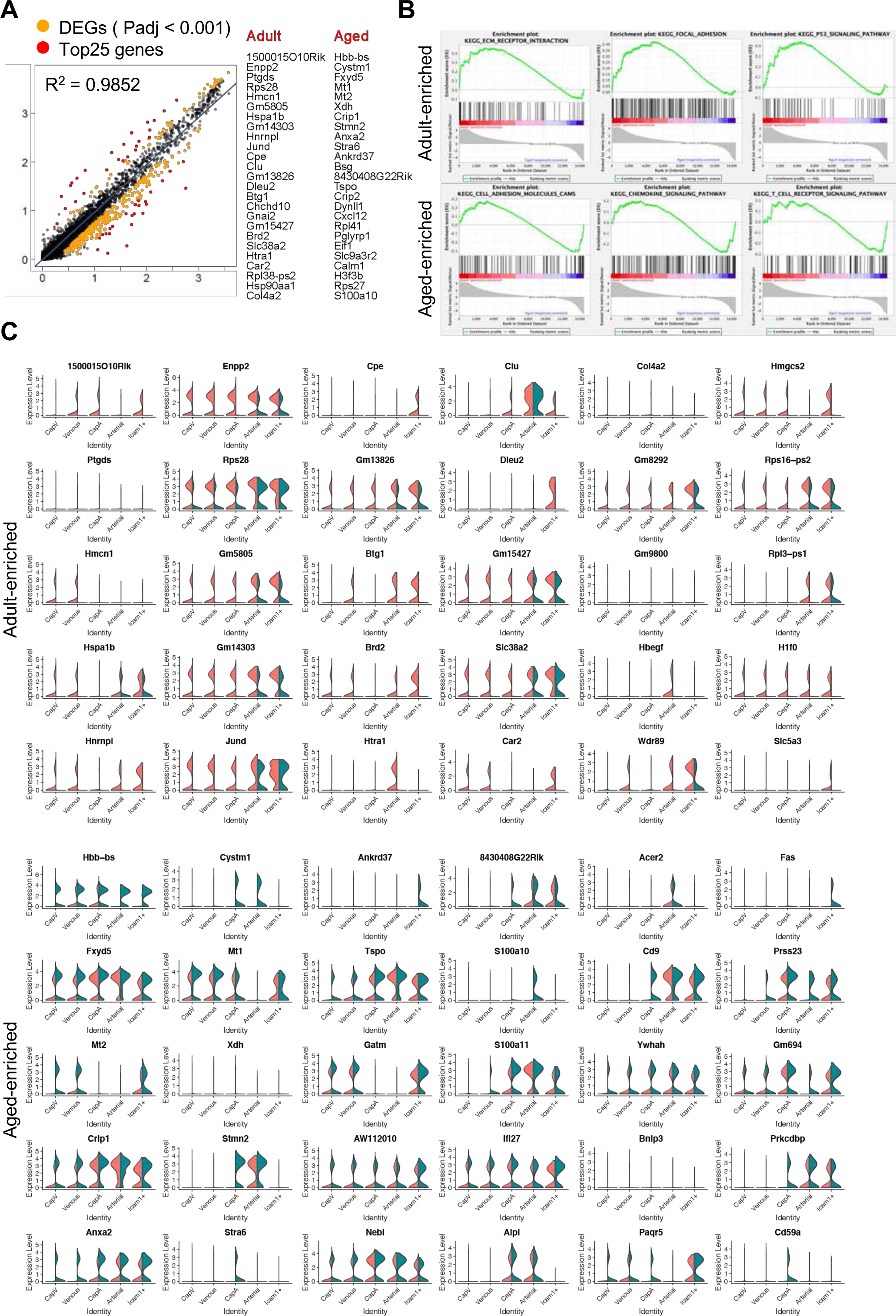
Differential gene expression analysis of adult and aged ECs. **(A)** Scatter dot plot of differentially expressed genes (yellow dots, Padj < 0.001) between ECs from adult and aged samples (407 genes). The 25 most significant genes for each sample (red dots) are listed. **(B)** Representative GSEA plots for overrepresented KEGG pathway gene sets in adult or aged ECs. **(C)** Violin plots of the top 30 differentially expressed genes in adult (red) and aged ECs (green).

**Figure 2-figure supplement 3.**
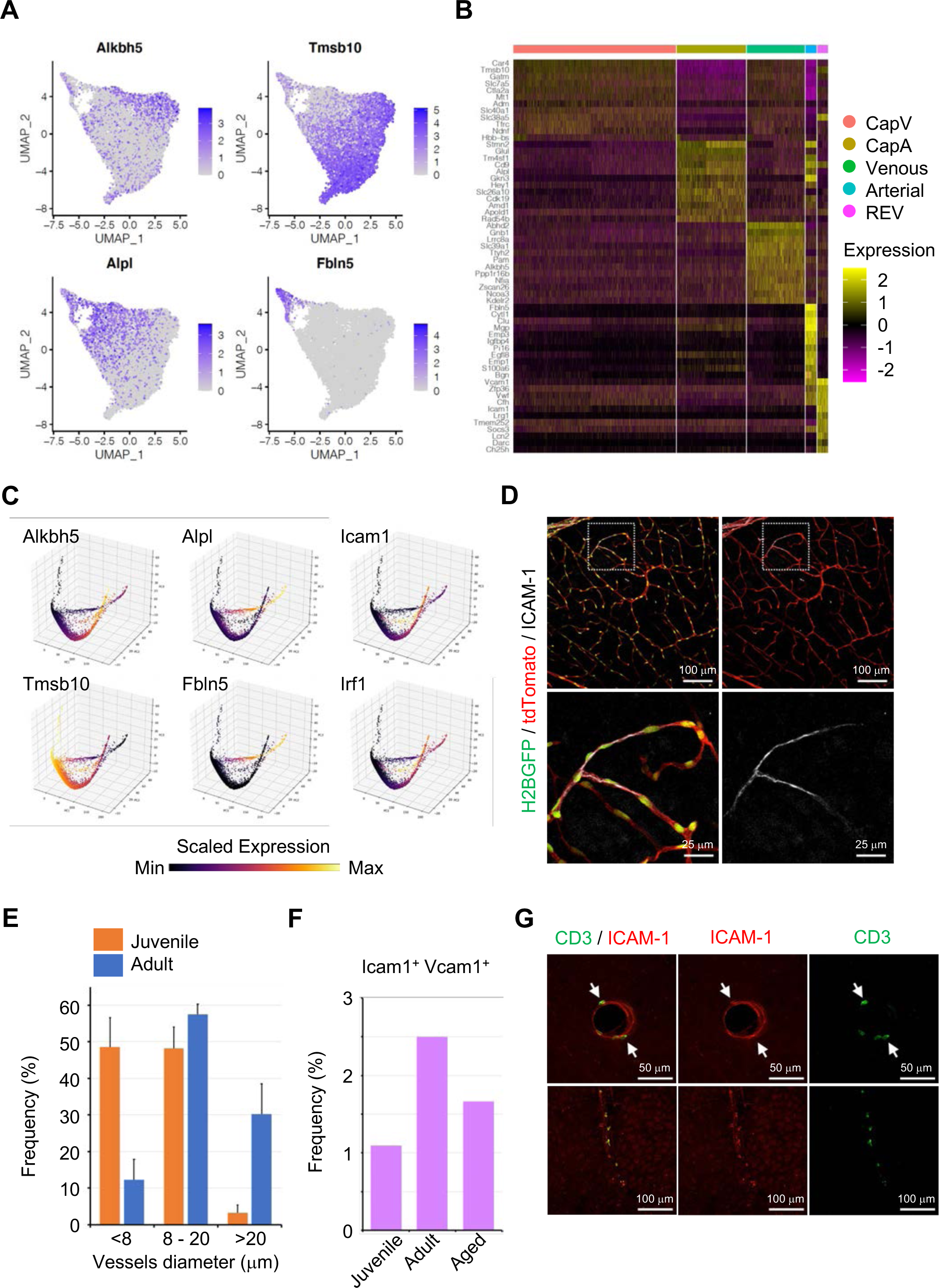
Subclustering analysis of ECs during homeostasis. **(A)** UMAP plots depicting the expression of representative marker genes for each sub-cluster: *Alkbh5* (Venous)*, Tmsb10* (CapV), *Alpl* (CapA) and *Fbln5* (Arterial). Colour key indicates scaled expression. **(B)** Heatmap of the top 12 marker genes for each sub-cluster of homeostatic ECs. **(C)** 3D PCA plots generated by MAGIC depicting the expression of selected subtype marker genes. Colour represents the scaled expression. **(D)** ICAM-1 immunostaining in P10 juvenile *Cdh5-H2BGFP/tdTomato* brain cortex. Scale bars, 100μm (top), 25μm (bottom). **(E)** Quantification of ICAM-1^+^ vessel diameter in brain cortical parenchyma. Error bars represent mean ± s.e.m. from three animals. **(F)** Frequency of *Icam1*/*Vcam1* double positive ECs among all ECs at different ages, according to the single cell RNA-seq data. **(G)** Immunofluorescence image showing CD3^+^ lymphocytes and ICAM-1^+^ vessels in the cerebral cortex 2 hr after LPS injection. Scale bar, 50μm (top), 100 μm (bottom).

**Figure 3-figure supplement 1.**
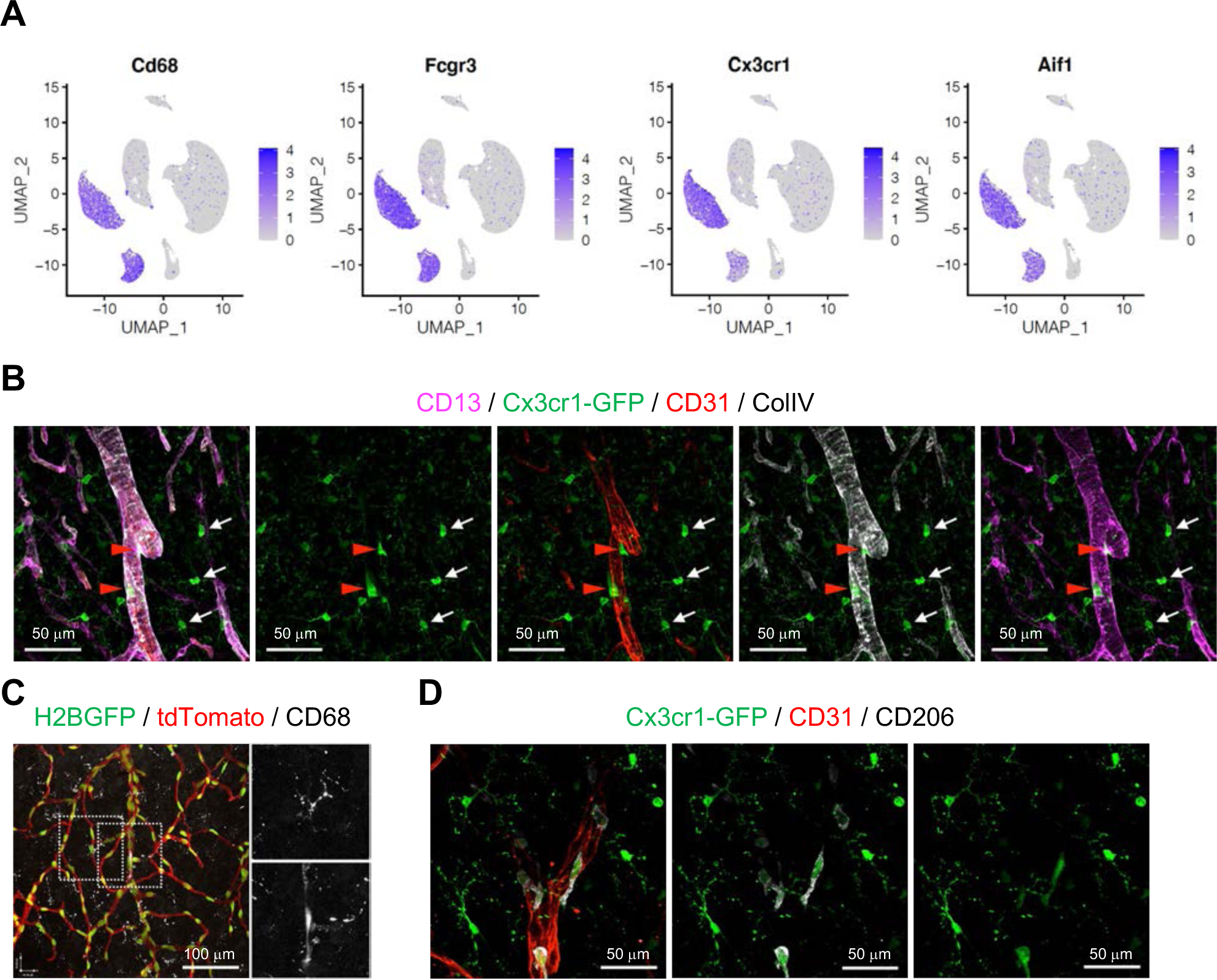
Distinct populations of microglia and PVMs. **(A)** UMAP plots depicting the expression of the known myeloid cell markers *Cd68, Fcgr3, Cx3cr1* and *Aif1*. Colour key represents the scaled expression. **(B)** Immunostaining showing CD13, GFP (*Cx3cr1-GFP*), CD31, and Collagen type IV (ColIV) in adult cortex. Scale bars, 50μm. **(C)** CD68 immunostaining in *Cdh5-H2BGFP/tdTomato* mouse brain cortex. Scale bar, 100μm. **(D)** *Cx3cr1-GFP* expression, CD31 and CD206 in sections of adult cortex. Scale bars, 50μm.

**Figure 4-figure supplement 1.**
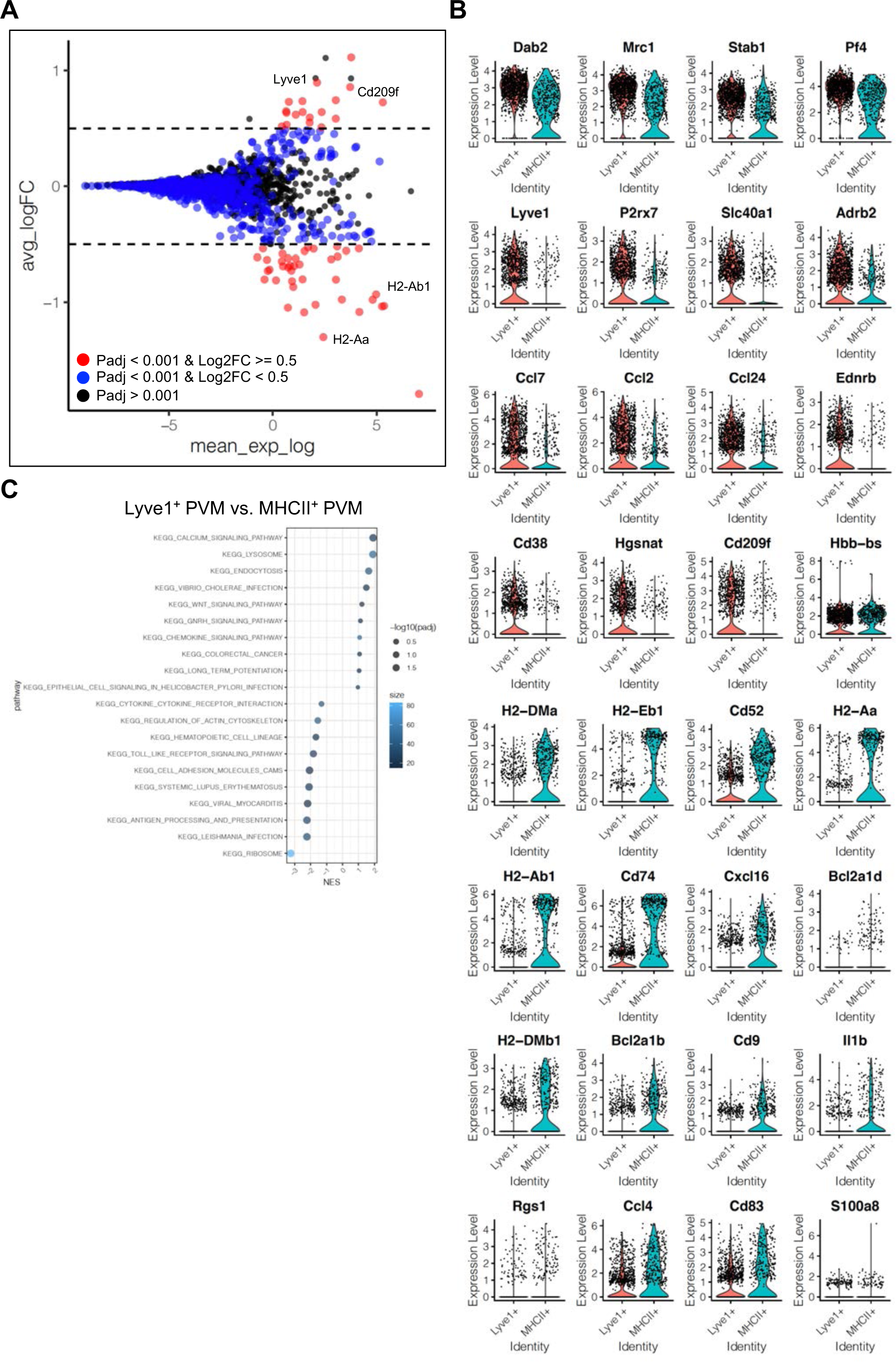
Differentially expressed genes in PVM subtypes. **(A)** MA plot of differentially expressed genes between Lyve1^+^ and MHCII^+^ PVM subtypes. Blue dots, p-adjusted value < 0.001; Red dots, p-adjusted value < 0.001 and Log2 fold change > 0.5. **(B)** Violin plots of the top differentially expressed genes between Lyve1^+^ and MHCII^+^ PVM subtypes. **(C)** Dot plot depicting top 10 enriched KEGG signalling pathways in GSEA for Lyve1^+^ and MHCII^+^ PVM subtypes. Dot size represents -log10 adjusted P values and colours indicate size of gene set.

**Figure 5-figure supplement 1.**
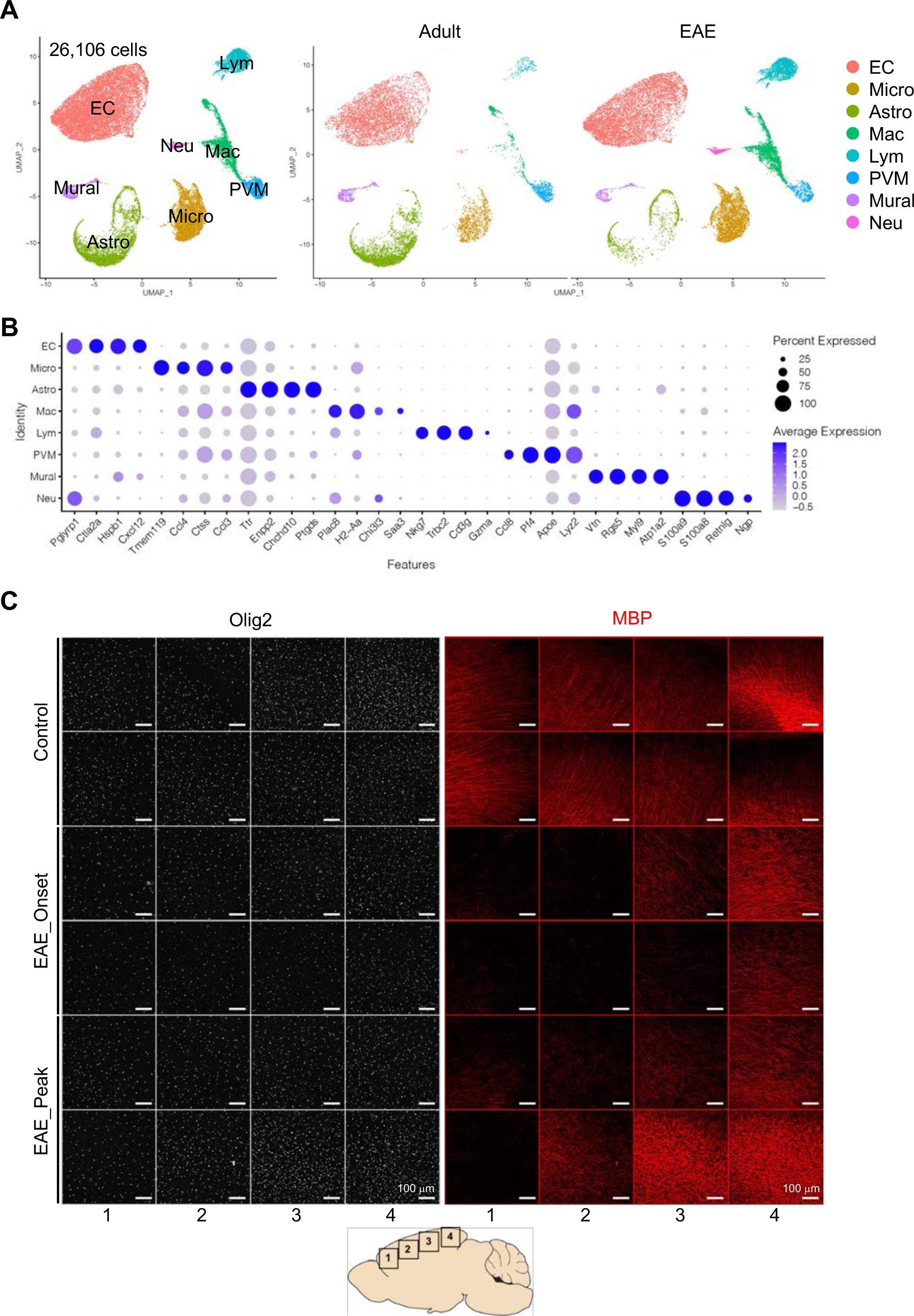
Single cell RNA-seq analysis of EAE mice. **(A)** UMAP plots of 26,106 myelin-depleted cells from adult and EAE cerebral cortex. Colours represent each cell type. **(B)** Dot plot displaying the top cell type-specific genes. Dot size represents the percentage of cells expressing the gene and colours reflect average expression of the gene. **(C)** Olig2 and MBP immunostaining in different regions (1-4) of brain cortex from control, onset and peak of EAE, as indicated. Each row shows an individual mouse. Scale bars, 100μm.

**Figure 5-figure supplement 2.**
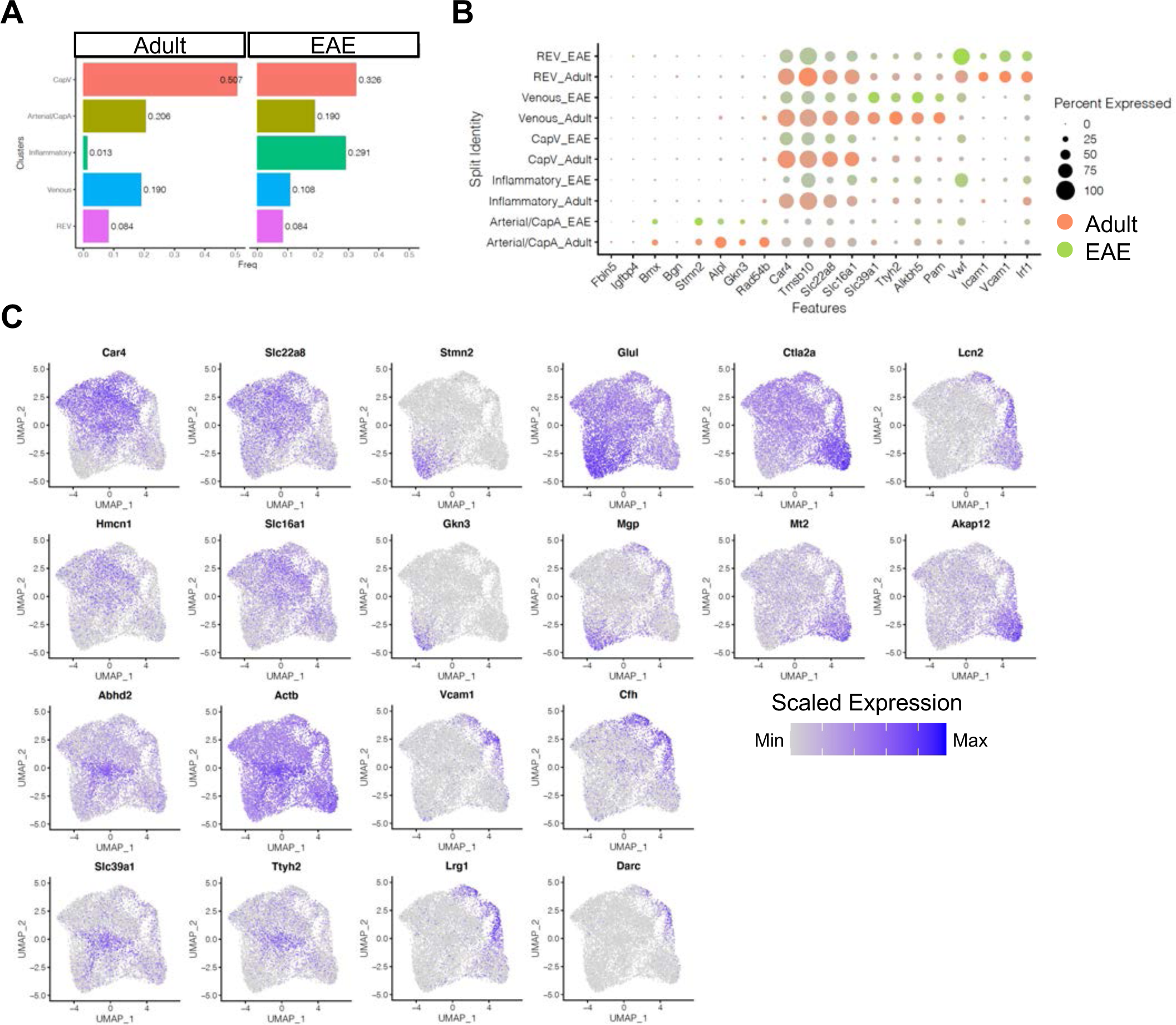
Subclustering of ECs from adult and EAE mice. **(A)** Bar plots indicating the frequency of EC subtypes in healthy adult and EAE brain. **(B)** Dot plot showing the expression of top sub-cluster-specific genes, with the dot size representing the percentage of cells expressing the gene and colours representing the average expression of the gene within a cluster. **(C)** UMAP plots depicting the expression of top marker genes for each EC sub-cluster. Colour represents the scaled expression.

**Figure 5-figure supplement 3.**
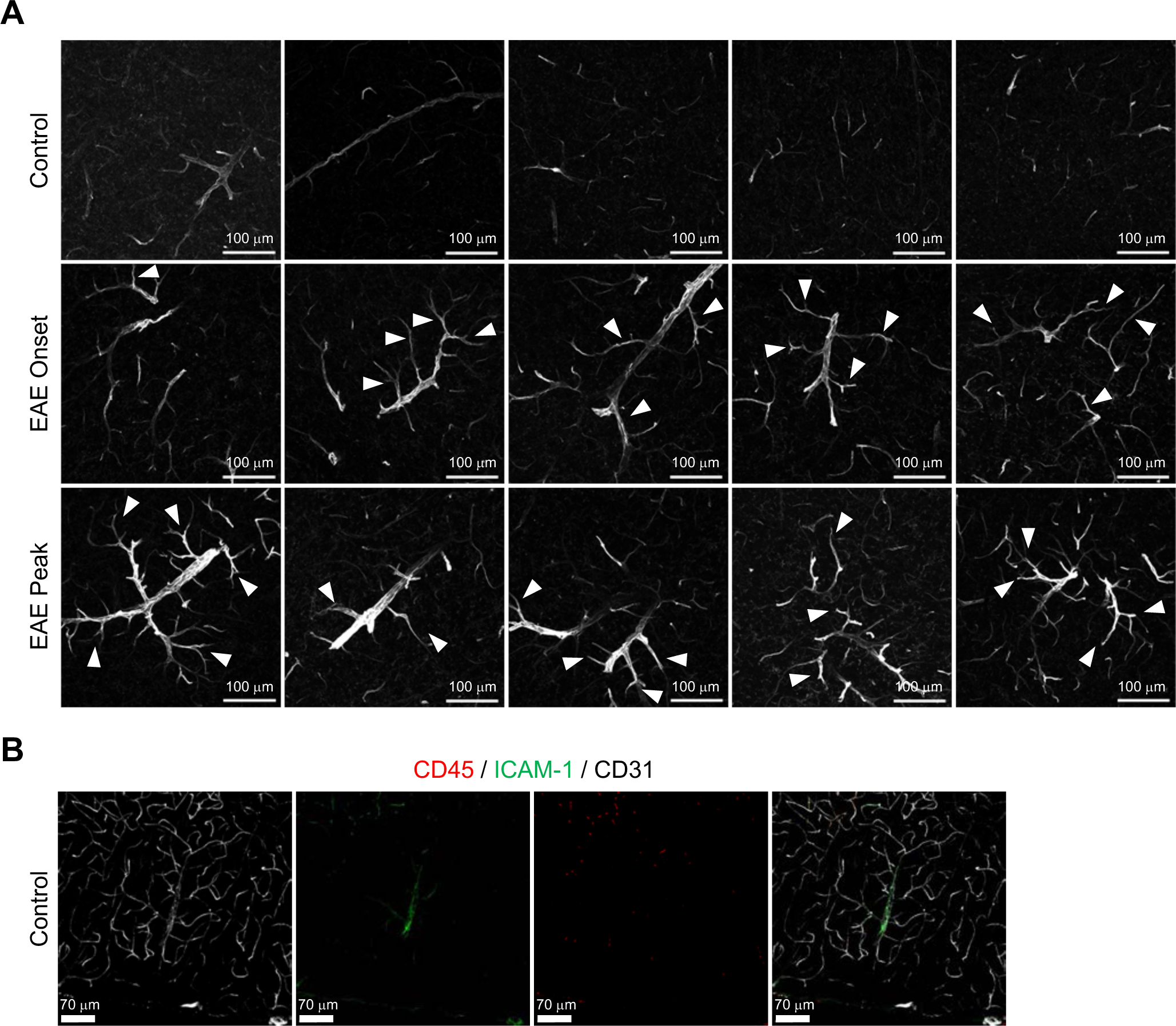
ICAM-1 expression in brain cortical parenchyma upon EAE. **(A)** Confocal images showing ICAM-1 immunofluorescence in cortex of control, EAE onset and peak mice. Arrowheads indicate expansion of ICAM-1 staining from REVs into adjacent venules and capillaries. Scale bars, 100μm. **(B)** Immunofluorescence images showing CD45, ICAM-1 and CD31 in the control adult cerebral cortex. Scale bars, 70μm.

**Figure 5-figure supplement 4.**
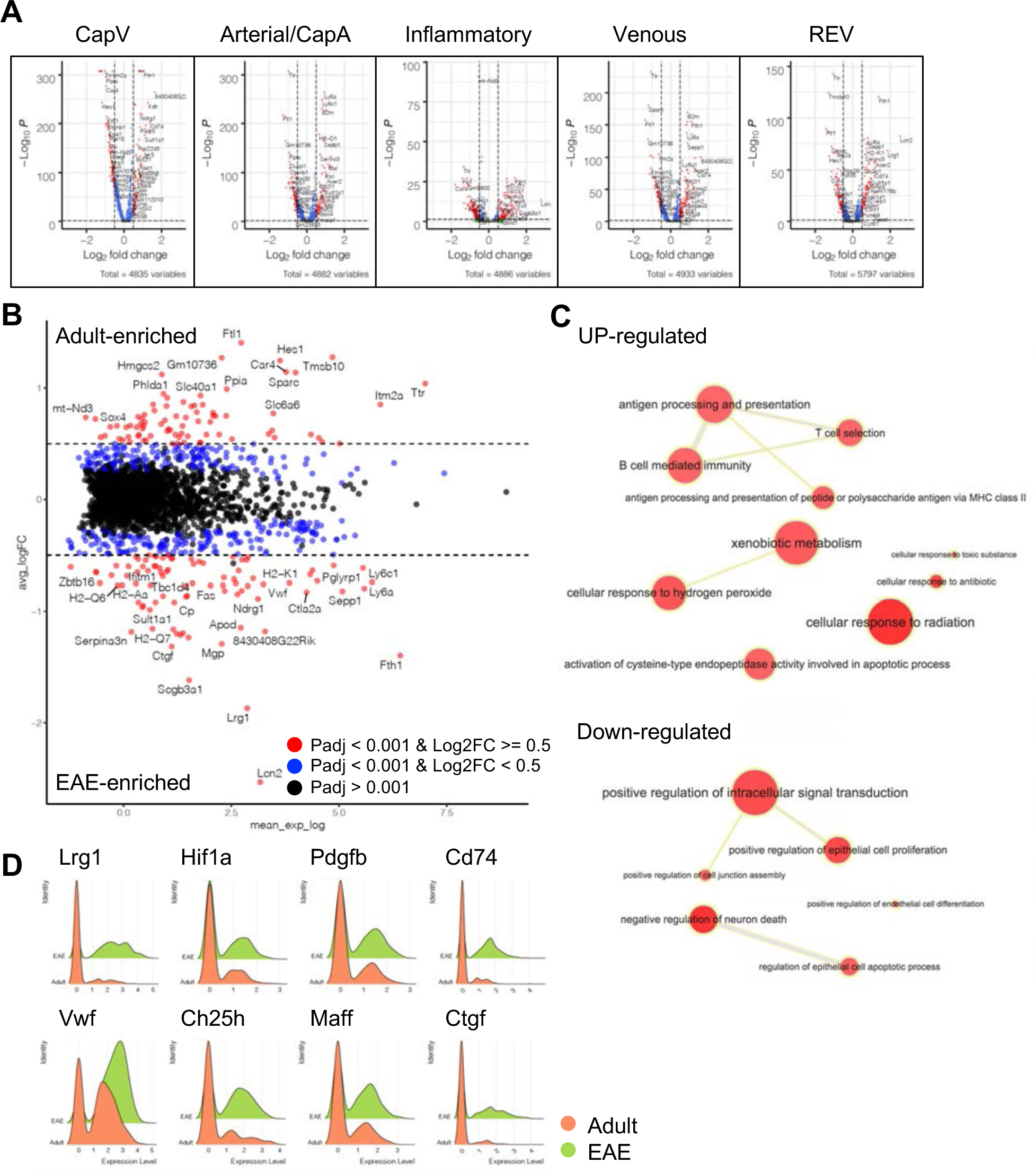
Differentially expressed genes between adult and EAE ECs. **(A)** Volcano plots depicting differentially expressed genes between adult and EAE EC subtypes. Blue dots, p-adjusted value < 0.001; Red dots, p-adjusted value < 0.001 and Log2 fold change > 0.5. (**B**) MA plot of differentially expressed genes between adult and EAE REVs. Blue dots, p-adjusted value < 0.001; Red dots, p-adjusted value < 0.001 and Log2 fold change > 0.5. (**C**) Significantly enriched GO terms for upregulated or downregulated genes in REV by EAE. Graph visualized by REVIGO. **(D)** Ridge plots of EAE-induced genes *Lrg1*, *Hif1a*, *Pdgfb*, *Cd74*, *Vwf, Ch25h, Maff* and *Ctgf* in REVs.

**Figure 6-figure supplement 1.**
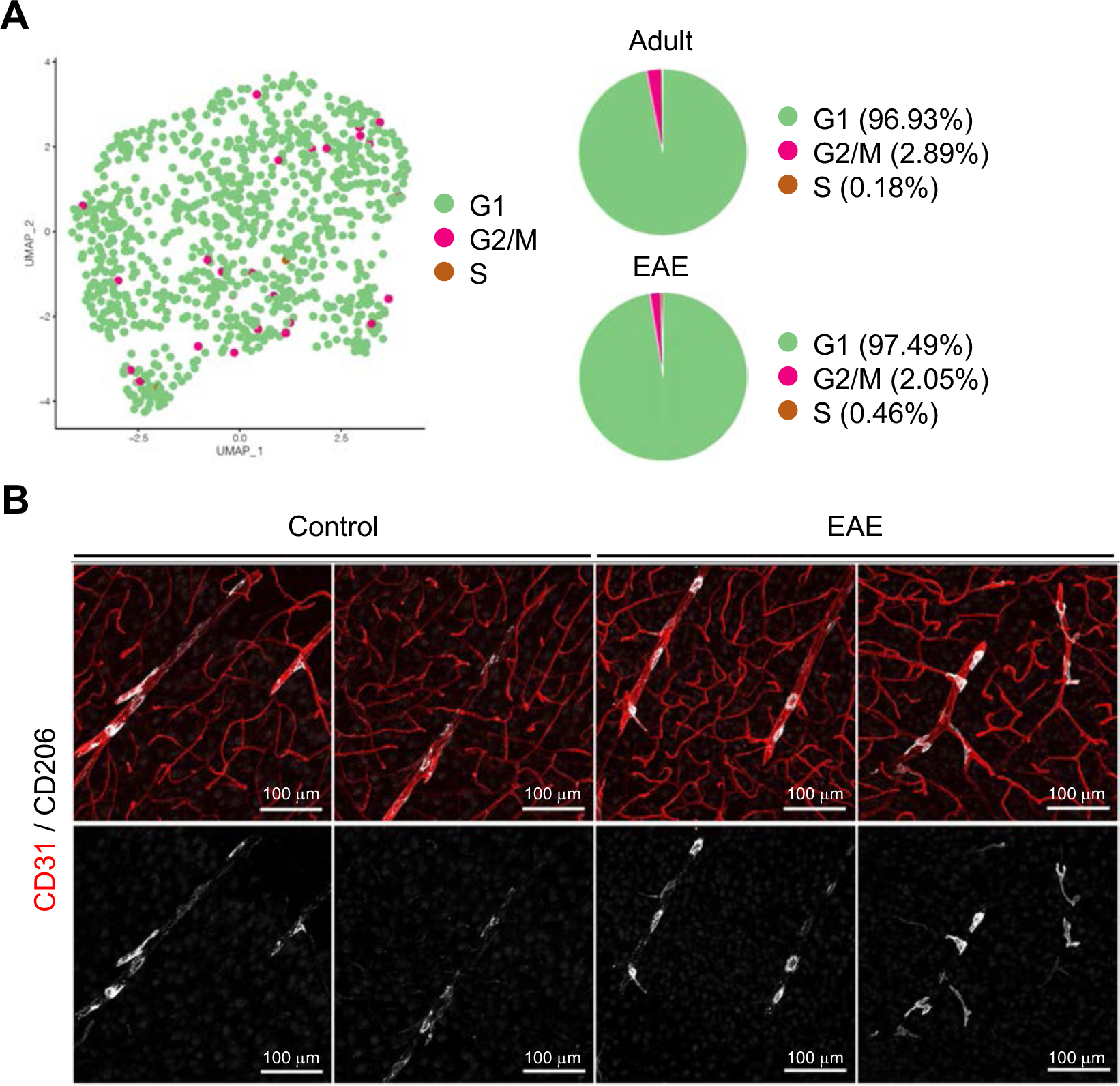
Cell cycle phases and localization of PVMs in EAE. **(A)** UMAP plot and pie chart of PVMs in adult and EAE brain cortex in the indicated cell cycle phases. **(B)** CD31 and CD206 immunostaining of brain cortex from control and EAE mice. Scale bars, 100μm.

**Figure 7-figure supplement 1.**
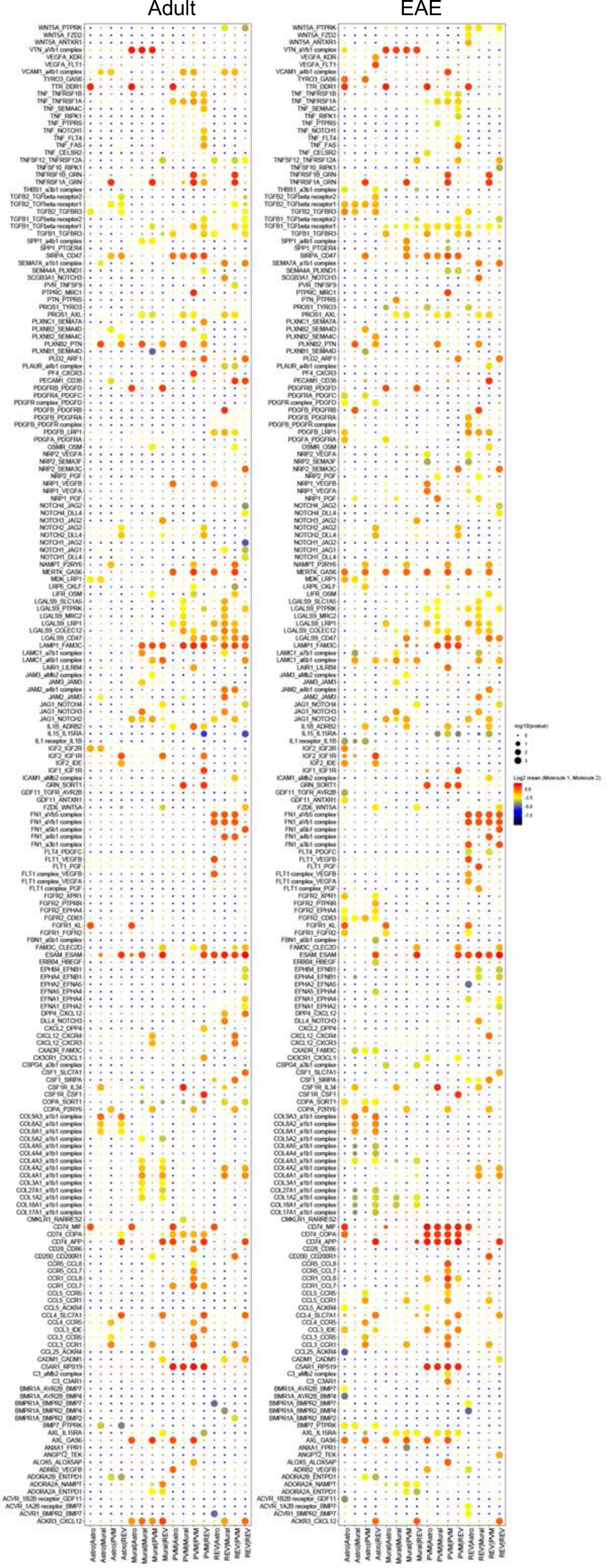
Interactome of BBB cell types in postcapillary venules. Overview of differential ligand-receptor interactions between REV EC, PVM, mural cell (Mural) and astroependymal cells (Astro) populations in adult control and EAE brain. Circle size indicates P values. Colour indicates mean average expression levels of interacting molecules.

**Figure 7-figure supplement 2.**
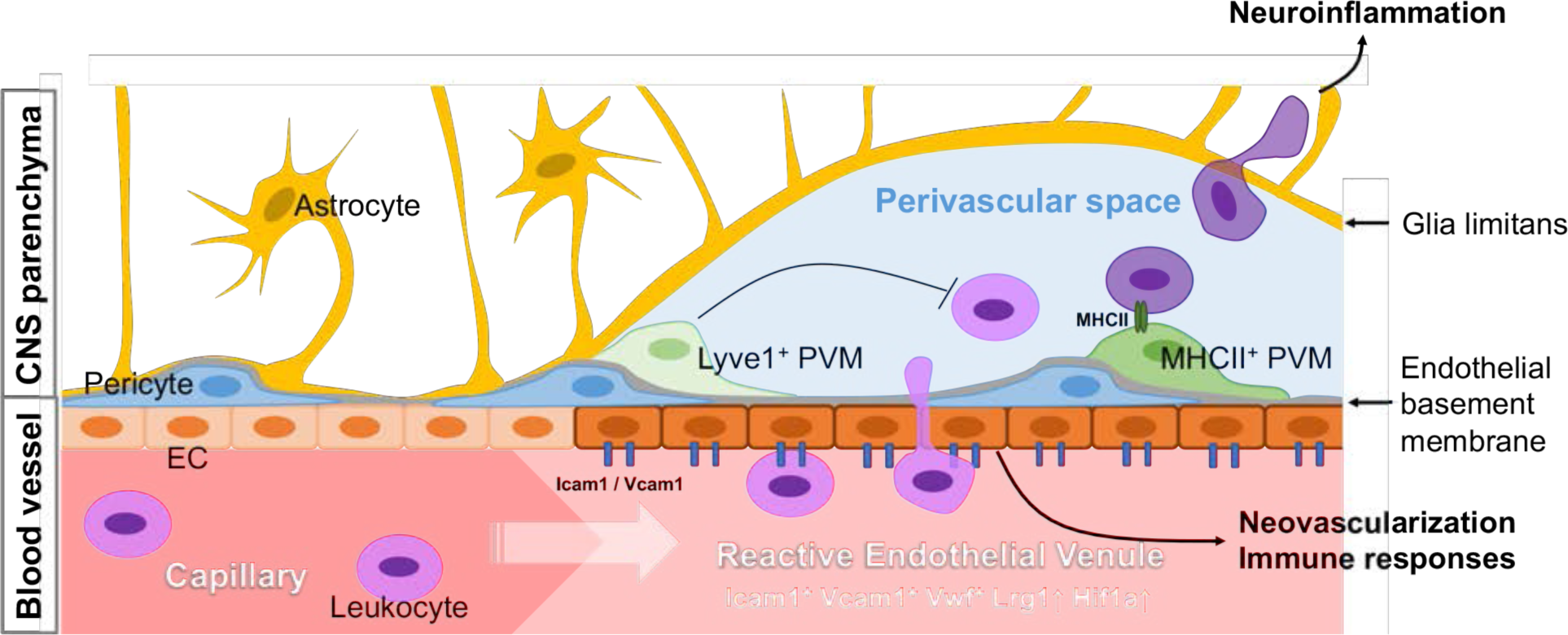
Schematic summary of experimental findings. Schematic summary of proposed findings. REVs are specialized cellular gateways in the CNS vasculature with associated PVMs as gatekeepers in the perivascular space, regulating the infiltration of circulating leukocytes into the brain parenchyma and vascular integrity during homeostasis and neuroinflammation.

